# Multiplexed Sequential Imaging in Living Cells with Orthogonal Fluorogenic RNA Aptamer/Dye Pairs

**DOI:** 10.1101/2023.04.20.537750

**Authors:** Ru Zheng, Rigumula Wu, Yuanchang Liu, Zhining Sun, Yousef Bagheri, Zhaolin Xue, Lan Mi, Qian Tian, Raymond Pho, Sidrat Siddiqui, Kewei Ren, Mingxu You

**Author notes:** These authors contribute equally (Ru Zheng, Rigumula Wu).

## Abstract

Single-cell detection of multiple target analytes is an important goal in cell biology. However, due to the spectral overlap of common fluorophores, multiplexed fluorescence imaging beyond two-to-three targets inside living cells remains a technical challenge. Herein, we introduce a multiplexed imaging strategy that enables live-cell target detection via sequential rounds of imaging-and-stripping process, which is named as “sequential Fluorogenic RNA Imaging-Enabled Sensor” (seqFRIES). In seqFRIES, multiple orthogonal fluorogenic RNA aptamers are genetically encoded inside cells, and then the corresponding cell membrane permeable dye molecules are added, imaged, and rapidly removed in consecutive detection cycles. As a proof-of-concept, we have identified in this study five *in vitro* orthogonal fluorogenic RNA aptamer/dye pairs (>10-fold higher fluorescence signals), four of which can be used for highly orthogonal and multiplexed imaging in living bacterial and mammalian cells. After further optimizing the cellular fluorescence activation and deactivation kinetics of these RNA/dye pairs, the whole four-color semi-quantitative seqFRIES process can now be completed in ∼20 min. Meanwhile, seqFRIES-mediated simultaneous detection of two critical signaling molecules, guanosine tetraphosphate and cyclic diguanylate, was also achieved within individual living cells. We expect our validation of this new seqFRIES concept here will facilitate the further development and potential broad usage of these orthogonal fluorogenic RNA/dye pairs for highly multiplexed and dynamic cellular imaging and cell biology studies.

## Introduction

Cellular processes are often regulated by a group of interactive biomolecules. The cellular distributions and dynamic correlations of these biomolecules play important roles in determining cell organization and the efficiency of cell signaling network^1–3^. The ability to analyze multiple biomolecules of interest, especially *in situ* inside living cells, is critical for understanding the molecular mechanisms and distinct signatures of both healthy and diseased cells. Fluorescence microscopy remains one of the most prominent and popular tools for the live-cell imaging of different target analytes. However, the broad fluorescence spectrum and large spectral overlap among common fluorophores have dramatically hindered their capability for the simultaneous detection of multiple target biomolecules inside living cells^4–6^. As a result, live-cell fluorescence imaging beyond two or three target analytes remains a common technical challenge. Although many efforts have been made towards multiplexed cellular detection, especially with recent development of vibrational^7–9^ and spectral imaging techniques^10,11^, as well as far-red fluorescent probes^12^, there is still an urgent need for simple and broadly applicable techniques, which can image multiple biomolecules in living systems just based on regular fluorescence microscopes available in a typical life science laboratory.

In fixed and membrane-permeabilized cell and tissue samples, the problem of fluorescence spectral overlap has been elegantly solved via sequential imaging approaches, such as MERFISH, seqFISH, etc^13–15^. In these techniques, target detection is achieved via dye-labeled DNA-or antibody-based fluorescent probes. Instead of imaging all the probes at a time, multiple rounds of labeling-imaging-and-stripping processes are conducted to allow highly multiplexed target detection. However, due to the requirement of cell permeabilization and fixation for the delivery and removal of these DNA/antibody-based fluorescent probes, current sequential multiplexing techniques are not capable for live-cell imaging or multiplexed molecular profiling in their native cellular context.

Herein, we report a novel sequential imaging platform that can be used for the live-cell multiplexed detection of different target analytes. This system is named <u>seq</u>uential <u>F</u>luorogenic <u>R</u>NA Imaging-<u>E</u>nabled <u>S</u>ensor (**seqFRIES**). Fluorogenic RNAs (**FR**) are single-stranded RNA aptamers that can specifically bind with otherwise non-fluorescent small-molecule dyes and activate their fluorescence signals^16^. One unique feature of the FR/dye system is that the binding between RNA aptamers and dyes is highly reversible and non-covalent (Fig. 1a). As a result, while the FR aptamers can be genetically encoded inside the cells, the corresponding membrane-permeable dye molecules can be externally synthesized, added to the cells for imaging, and then easily washed away to turn off fluorescence signals. Sequential multiplexed detection in living cells can thus be possibly achieved via rounds of rapid imaging-and-stripping processes using this non-covalent fluorogenic RNA/dye-based system (Fig. 1b).

**Fig. 1.**
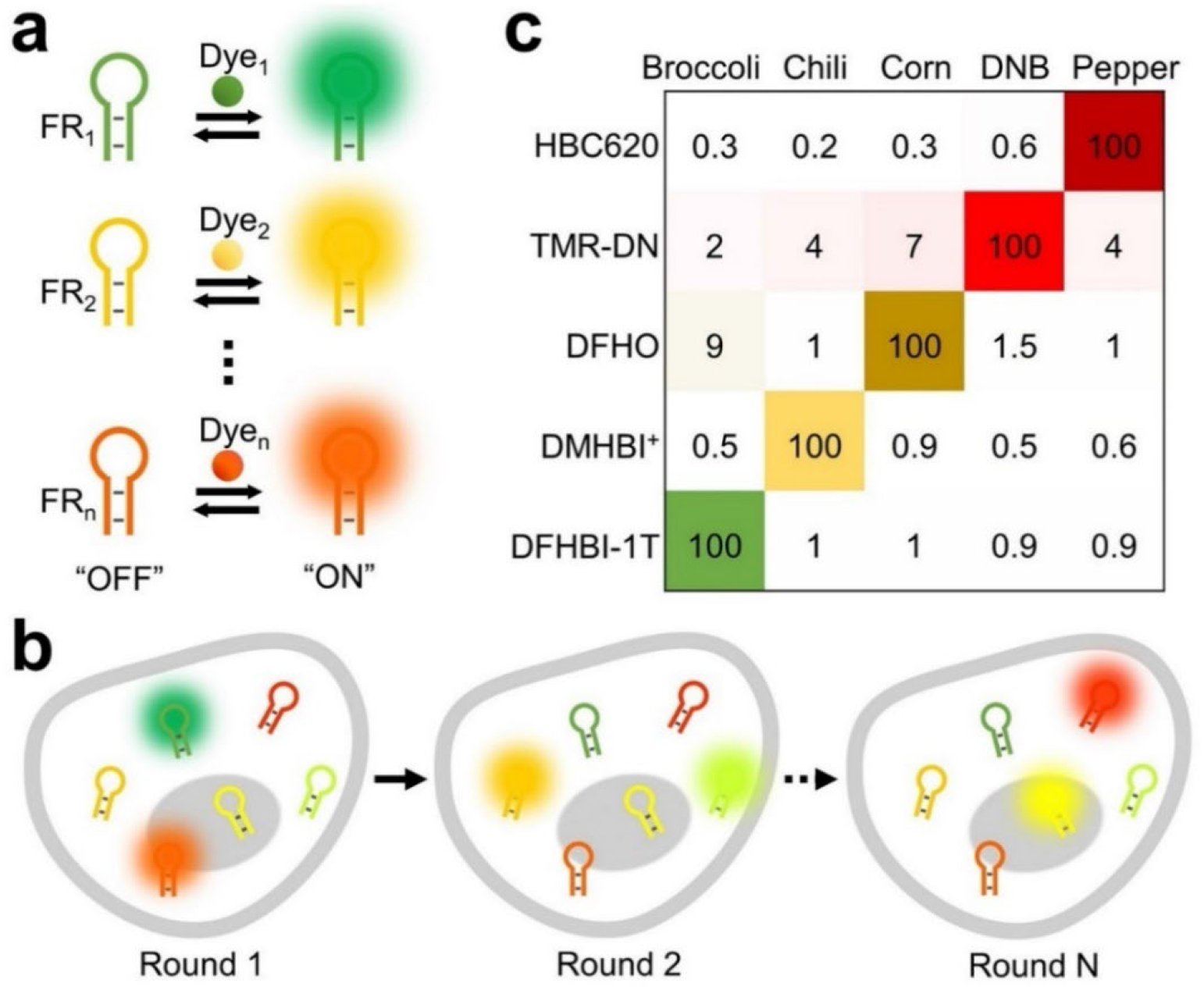
Working principle of fluorogenic RNA (FR) and sequential multiplexed imaging system. **a** Upon specific reversible binding, orthogonal FR/dye pairs can turn on their fluorescence signals independent from each other. **b** After genetically encoding orthogonal FRs in living cells, sequential multiplexed detection can be achieved via rounds of imaging-and-stripping process by adding and removing the cognate membrane-permeable small-molecule dyes. **c** *In vitro* characterization of five identified orthogonal FR/dye pairs. More experimental details are shown in Figure S1. Both the number and brightness of each block indicate the intensity of mean fluorescence signals as measured from at least three independent replicates.

To develop such a seqFRIES platform, the identification of orthogonal FR/dye pairs with fast cellular fluorescence activation/deactivation kinetics is highly critical. This is because the potential multiplexing capability of seqFRIES will be determined by the number of FR/dye pairs that can function independently; while the overall imaging efficiency and throughput will be controlled by the time required for each round of labeling, imaging, and stripping. In this study, we first identified the largest set of orthogonal fluorogenic RNA/dye pairs reported so far, including five FR/dye pairs that can function independently *in vitro*, four pairs of which can be used for multiplexed sequential imaging in living bacterial and mammalian cells. We have further optimized the cellular fluorescence turn-on and turn-off rate for each of these four FR/dye pairs. Under optimized imaging-and-stripping conditions, semi-quantitative four-color target detection in living cells can be achieved within ∼20 min.

In addition, these FR aptamers can be modularly engineered into sensors for the cellular recognition and detection of different target analytes^17–19^. Using metabolites and signaling molecules as sample targets, our results have also demonstrated the potential usage of seqFRIES for studying the correlation between cellular distributions of different biomolecules in single living cells. By adapting a new concept of live-cell sequential imaging, we expect the development of this seqFRIES platform will open the door for new applications in the field of live-cell multiplexed imaging and single-cell molecular profiling, which can be performed with conventional fluorescence microscopes.

## Results

### *In vitro* identification of orthogonal FR/dye pairs

Our first goal was to test if some existing fluorogenic RNA aptamers and their cognate dyes can be used as orthogonal pairs for potential multiplexed detection. We selected six previously reported FR/dye pairs, including Broccoli/DFHBI-1T, Chili/DMHBI^+^, Corn/DFHO, DNB/TMR-DN, Mango II/YO3-biotin, and Pepper/HBC620 (the RNA sequences are listed in Supplementary Table 1 and the chemical structures of the dyes are shown in Supplementary Fig. 1)^20–28^. After *in vitro* transcription of six fluorogenic RNA aptamers and respectively mixed with each of the six dyes, the fluorescence spectra of all these 36 FR/dye pairs were then measured in solutions containing 1–5 μM fluorogenic RNA and 0.5–20 μM dye molecules. The concentration range of these RNAs and dyes was chosen based on the recommended conditions in the original reports^20–28^. Under optimized conditions, the DFHBI-1T, DFHO, DMHBI^+^, HBC620, and TMR-DN dyes exhibited great orthogonality. These dye molecules can only be activated by their corresponding fluorogenic RNAs, with at least 10–600-fold higher fluorescence signals than any other tested RNA aptamers (Fig. 1c and Supplementary Fig. 1). In contrast, the YO3-biotin dye can be nonspecifically activated by almost all the attempted FR aptamers and thus was not used in the following studies.

We next evaluated if the fluorescence of each FR/dye conjugate will be influenced by the presence of other RNA aptamers in the same solution. By varying the concentrations of these RNA aptamers in the range of 1–12 μM, linearly increased fluorescence signals were observed in all types of solutions containing either individual FR/dye pair or a mixture of RNAs with each dye molecule (Supplementary Fig. 2). Such a linear correlation can be critical for potential quantitative detection^29^. Meanwhile, our data indicated that the Broccoli/DFHBI-1T and Pepper/HBC620 fluorescence intensities were not affected by other RNA aptamers, while a slight decrease (∼28–46%) in the DNB/TMR-DN and Chili/DMHBI^+^ fluorescence signals were shown in the RNA mixture, and interestingly, Corn/DFHO signals were even enhanced by ∼30% in the mixture. It is possible that some RNA–RNA interactions may exist in these mixtures, but still these five FR/dye pairs, i.e., Broccoli/DFHBI-1T, Chili/DMHBI^+^, Corn/DFHO, DNB/TMR-DN, and Pepper/HBC620, can function orthogonally *in vitro* and were identified as the potential candidates for developing the multiplexed seqFRIES system.

### Cellular fluorescence activation and deactivation kinetics of the FR/dye pairs

We next cloned each FR aptamer respectively into a pETDuet vector and then the vector was transformed into BL21 Star (DE3) *E. coli* cells. Upon adding each cognate dye, strong cellular fluorescence signals were clearly observed in the case of Broccoli/DFHBI-1T, Corn/DFHO, DNB/TMR-DN, and Pepper/HBC620 (Supplementary Fig. 3a). However, the addition of DMHBI^+^ failed to induce any fluorescence signals in Chili-expressing cells. We speculate that the DMHBI^+^ dye may have poor cell membrane permeability, and in fact, the Chili/DMHBI^+^ pair has never been applied for cellular imaging before. As a consequence, only the other four FR/dye pairs were selected for later investigations.

Considering the importance of fast imaging-and-stripping process in sequential imaging techniques, we next wanted to assess the cellular fluorescence activation and deactivation kinetics of each FR/dye pair. As shown in Supplementary Fig. 3b, c, after adding the DFHO, DFHBI-1T, and HBC620 dyes into the corresponding Corn-, Broccoli- or Pepper-expressing *E. coli* cells, very fast cellular fluorescence enhancement was observed: ∼80% of maximal cellular signals were reached within ∼7 min, 8 min, and 13 min, respectively. While a slower fluorescence activation was exhibited for the DNB/TMR-DN pair, ∼45% of the maximal cellular fluorescence could be detected in ∼30 min. It is worth noting that because of the quite high brightness of DNB/TMR-DN, their cellular fluorescence signals can still be clearly visualized even at <10% of the maximal level.

We also measured how quickly the cellular fluorescence signals can be stripped upon removing the dye molecules. In these experiments, FR-expressing *E. coli* cells were first incubated with the corresponding dye molecule for 50 min to reach near-maximum fluorescence intensities. Then the medium was swapped into a dye-free Dulbecco’s phosphate-buffered saline (DPBS) for two rounds of 1-min incubation. Consistent with the fluorescence activation results, cellular fluorescence from the Broccoli/DFHBI-1T, Corn/DFHO, and Pepper/HBC620 pairs could be easily removed: after the total of 2-min washing, only ∼17%, 20%, and 17% signals remained in these bacterial cells (Supplementary Fig. 4a, b). With very fast fluorescence activation and deactivation kinetics, these three FR/dye pairs can be ideally used for developing the seqFRIES sequential imaging platform.

In the case of DNB/TMR-DN, the same 2-min washing protocol is not enough to remove cellular fluorescence signals (Supplementary Fig. 4a, b). Even after further increasing the incubation time and rounds of washing, our results indicated that ∼45% and 25% of DNB/TMR-DN signals remained in the cells after two or five rounds of 6-min washing (Supplementary Fig. 4c, d). Apart from washing, we wondered if the stripping of cellular DNB/TMR-DN fluorescence signals can also be achieved via photobleaching or competitive binding. Indeed, after 1-min irradiation with a 1 mW/cm^2^ power 561 nm laser, cellular DNB/TMR-DN fluorescence was reduced to ∼1% (Supplementary Fig. 5). Similarly, by adding a non-fluorescent small-molecule ligand, DN-PEG3-amine, to compete with TMR-DN for the dinitroaniline binding site in the DNB aptamer, cellular DNB/TMR-DN signals can also be rapidly reduced to ∼15% of the original intensity within ∼5 min (Supplementary Fig. 5). All these data supported that the reversible binding between FRs and dyes can be potentially applied for achieving rapid imaging-and-stripping cycles.

### Sequential cellular imaging of multiple FR/dye pairs

Our next goal was to test if Broccoli/DFHBI-1T, Corn/DFHO, DNB/TMR-DN, and Pepper/HBC620 can be used to develop a multiplexed sequential imaging system. We first wanted to validate the orthogonality of these FR/dye pairs inside living cells. For this purpose, BL21 Star (DE3) *E. coli* cells that respectively express Broccoli, Corn, DNB, or Pepper RNAs were separately incubated with the DFHBI-1T, DFHO, TMR-DN, or HBC620 dye for 30 min. The fluorescence imaging results clearly revealed that each kind of dye molecule could only be specifically activated by the cognate RNA aptamer (Fig. 2a, b). Indeed, these four FR/dye pairs maintained their great orthogonality inside bacterial cells for potential multiplexed detection.

**Fig. 2.**
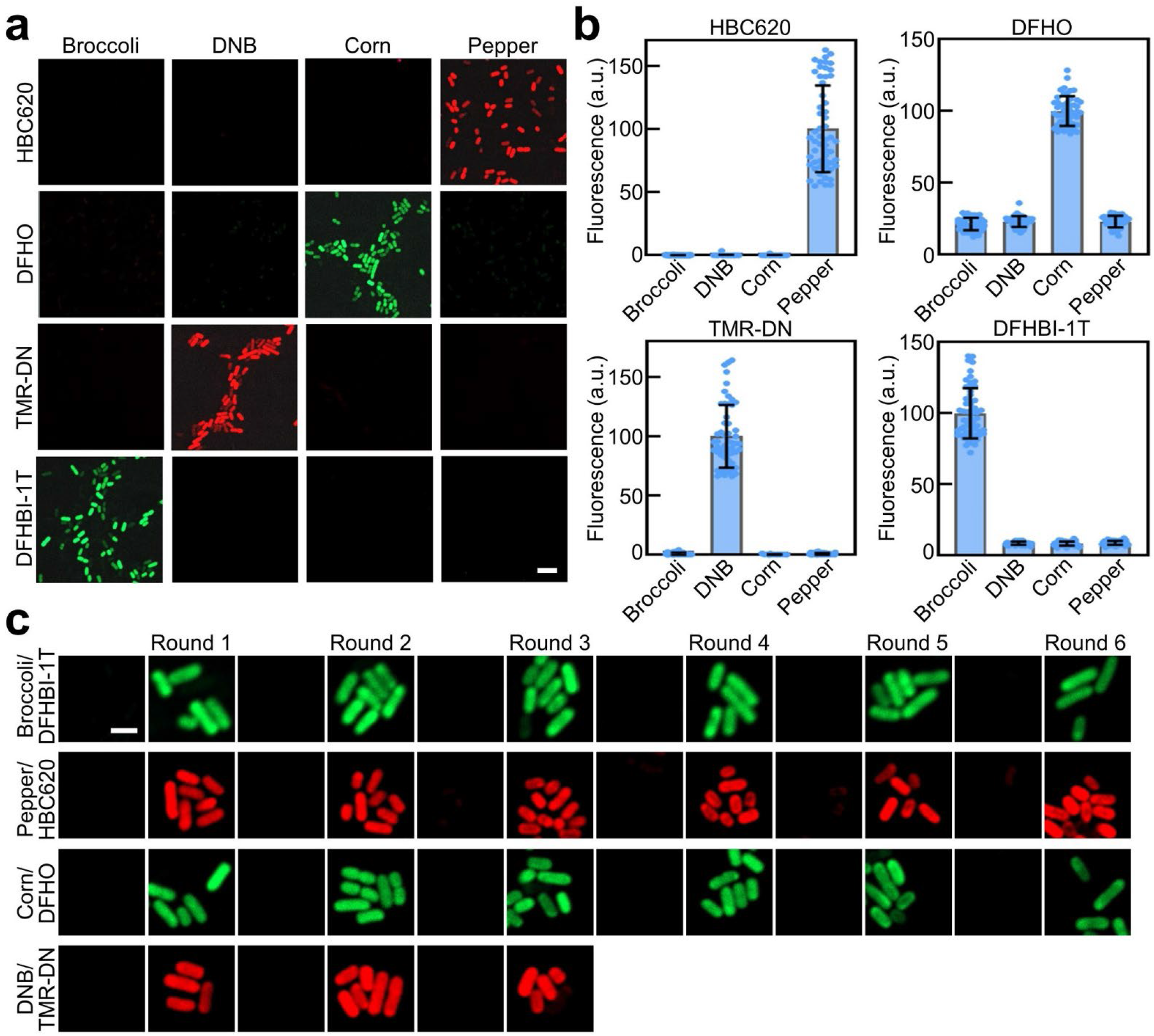
Orthogonality and sequential imaging feasibility of FR/dye pairs in live *E. coli* cells. **a** Fluorescence images were taken in BL21 Star (DE3) cells that express Broccoli, DNB, Corn, or Pepper RNA after a 30-min incubation with 200 μM DFHBI-1T, 1 μM TMR-DN, 10 μM DFHO, or 1 μM HBC620 dye molecules. Scale bar, 5 μm. **b** Cellular fluorescence intensities of the orthogonality experiments shown in panel (**a**) were also quantified in ∼50 individual cells in each case. Shown are the mean and standard deviation (SD) values from cellular images taken from at least three independent replicates. **c** In BL21 Star (DE3) cells that express all four FRs (FDB-FPC), fluorescence images were taken after each round of 30-min incubation with 200 μM DFHBI-1T, 1 μM TMR-DN, 10 μM DFHO, or 1 μM HBC620 dye molecules. Three times of 1-min DPBS washing in the case of DFHBI-1T, DFHO, and HBC620, or five times of 6-min washing for TMR-DN, were performed right after each round of imaging to strip cellular fluorescence. Scale bar, 5 μm.

We next prepared a plasmid construct that can simultaneously express all four fluorogenic RNAs within the same cells. A pETDuet vector was chosen here because it contains two expression cassettes. DNB and Broccoli were fused via a three-way junction F30 RNA scaffold and inserted into one cassette (Supplementary Fig. 6a). Similarly, Pepper and Corn were conjugated into another F30 scaffold and cassette to allow the co-expression of all four RNA aptamers using one plasmid construct, which is named **FDB-FPC**. The F30 scaffold is used here to improve the proper folding of RNA aptamers and to reduce the interference among RNA sequences^30^. Interestingly, while the Pepper signal was not changed after being inserted into the F30 scaffold, the fluorescence intensities of *in vitro* transcribed Broccoli, Corn, and DNB within these F30 scaffolds were increased by ∼0.6-, 0.7- and 1.8-fold compared to those without the scaffold (Supplementary Fig. 6b). This is likely due to the improved RNA folding in the F30 scaffold. After cloning this FDB-FPC plasmid into BL21 Star (DE3) cells, we incubated the cells with DFHBI-1T, DFHO, TMR-DN, and HBC620 dyes, respectively, for 30 min. Bright cellular fluorescence signals were observed in all these conditions (Supplementary Fig. 6c). In the presence of DFHO, the cellular fluorescence intensities were even 1.7-fold higher than cells that only express the Corn aptamer (Supplementary Fig. 6d).

Using these FDB-FPC-expressing *E. coli* cells, we next investigated if the fluorescence signals from each of these four FR/dye pairs can be repeatedly imaged and stripped. In the case of DFHBI-1T, DFHO, and HBC620, cellular fluorescence signals were measured after each round of 30-min incubation, and three times of 1-min washing were performed right after each round of imaging to strip cellular fluorescence signals (Fig. 2c). During a total of six rounds of imaging-and-stripping, >70% of original cellular fluorescence signals can still be observed for all the three FR/dye pairs of Broccoli/DFHBI-1T, Corn/DFHO, and Pepper/HBC620 (Supplementary Fig. 7). Such slightly reduced fluorescence intensities are likely due to the cellular degradation of fluorogenic RNAs during these over 3-hour-long experiments. In the case of DNB/TMR-DN conjugate, considering its relatively slow fluorescence activation and deactivation kinetics, we only tested three rounds of sequential imaging by adding TMR-DN for a 30-min incubation and then removing the dye via five times of 6-min washing. Similar to other FR/dye pairs, ∼100% of cellular DNB/TMR-DN signals remained observed during this alternate fluorescence activation and deactivation procedure (Fig. 2c and Supplementary Fig. 7). All these results together suggested the robustness of FR/dye fluorescence signals during the repeated imaging-and-stripping process.

To test the feasibility of these FR/dye pairs-based multiplexed imaging system, we next tried to sequentially image cellular Pepper/HBC620, Corn/DFHO, Broccoli/DFHBI-1T, and DNB/TMR-DN fluorescence signals, one at a time, in the above-mentioned BL21 Star (DE3) *E. coli* cells that express FDB-FPC (Fig. 3a). The HBC620, DFHO, DFHBI-1T, and TMR-DN dyes were added in order, and each incubated for 20 min before imaging. Between each round of detection, three times of brief washings (1–3 min) were performed to ensure the removal of the previously added dyes. Indeed, >90% of cellular fluorescence signals were stripped before the next round of imaging. Very nice correlations in the cellular fluorescence intensities (Pearson’s correlation coefficient r in the range of 0.79–0.91) were observed among all these four FR/dye pairs (Fig. 3b). Almost 100% of *E. coli* cells exhibited bright fluorescence signals during all four rounds of imaging cycles. These four FR/dye pairs can thus really be used to develop the proposed seqFRIES platform.

**Fig. 3.**
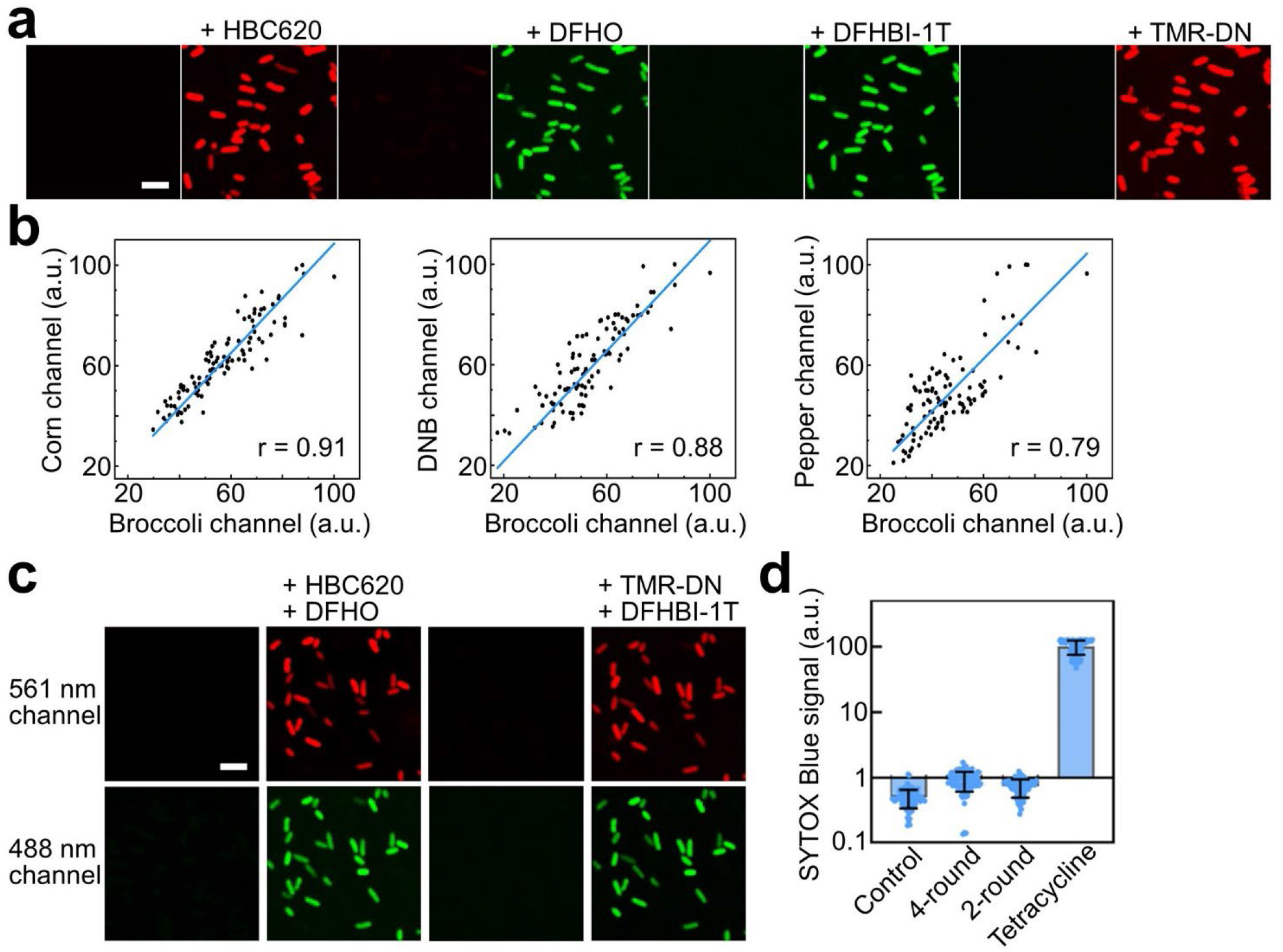
Sequential multiplexed imaging in living bacterial cells. **a** Fluorescence images were taken in BL21 Star (DE3) cells that express all four FRs (FDB-FPC) after a sequential 20-min incubation with 1 μM HBC620, 10 μM DFHO, 200 μM DFHBI-1T, or 1 μM TMR-DN dye molecules. Three times of 3-min DPBS washing for HBC620, and three times of 1-min washing in the case of DFHO and DFHBI-1T, were used respectively to strip cellular fluorescence. Scale bar, 5 μm. **b** Correlations of cellular Broccoli, Corn, DNB and Pepper fluorescence signals as measured within ∼100 individual cells after the 4-round imaging protocol shown in panel (**a**). Pearson’s correlation coefficient r was determined in each case where a value between 0.79 and 0.91 indicates a moderate-to-strong positive correlation. **c** Fluorescence images were taken in BL21 Star (DE3) cells that express all four FRs (FDB-FPC) after a rapid sequential 5-min incubation with a mixture of 1 μM HBC620 and 10 μM DFHO, or a mixture of 200 μM DFHBI-1T and 1 μM TMR-DN. Three times of 3-min DPBS washing for HBC620 and DFHO were performed to strip cellular fluorescence. Scale bar, 5 μm. **d** Cytotoxicity measurement was performed by incubating BL21 Star (DE3) cells with 1 μM SYTOX Blue for 5 min before (control) and after a 4-round (panel **a**) or 2-round (panel **c**) sequential imaging process. Cells that were pre-treated with 1 mM tetracycline for 30 min was used as the positive control. Cellular SYTOX Blue fluorescence signals were measured using ∼50 individual cells in each case upon a 405 nm laser irradiation. Shown are the mean and SD values from cellular images taken from at least three independent replicates.

We further studied if these four FR/dye pairs can be potentially imaged within a shorter time frame by simultaneously detecting two pairs in one imaging round. It is worth mentioning that due to significant spectral overlap, in our case, Broccoli/DFHBI-1T and Corn/DFHO cellular fluorescence were detected using the same filter set (525/50 nm) upon irradiation via a 488 nm laser. Similarly, Pepper/HBC620 and DNB/TMR-DN were both excited by a 561 nm laser, and the emissions were collected in the range of 575–625 nm. As a result, these FR/dye pairs of similar color cannot be distinguished or imaged together. Our previous work demonstrated that Broccoli/DFHBI-1T and DNB/TMR-DN exhibit a large difference in their excitation/emission spectra and can be simultaneously imaged in the cells^31^. We thus wondered whether Corn/DFHO and Pepper/HBC620 could also be imaged together. Indeed, minimal spectral overlap and almost no fluorescence interference was observed when the DFHO and HBC620 dyes were added to FDB-FPC-expressing BL21 Star (DE3) cells (Supplementary Fig. 8a). Corn/DFHO and Pepper/HBC620 signals could be clearly distinguished in each individual cell. Similarly, Broccoli/DFHBI-1T and DNB/TMR-DN fluorescence signals are also distinct from each other in these FDB-FPC-expressing cells (Supplementary Fig. 8a).

To achieve rapid sequential imaging, we decided to add DFHO and HBC620 together during the first round of detection, and then incubated a mixture of DFHBI-1T and TMR-DN dyes for the second round of imaging. Based on the cellular fluorescence activation kinetics data (Supplementary Fig. 3), we also decided to reduce the incubation time of each dye pair to 5 min. By performing three times of 3-min washing between two imaging rounds, the whole seqFRIES process can be completed within ∼20 min. Following this protocol, the fluorescence signals from all four FR/dye pairs can be clearly visualized within ∼100% of FDB-FPC-expressing cells (Fig. 3c), with nice correlations in the four-channel cellular fluorescence intensities (Pearson’s correlation coefficient r in the range of 0.68– 0.90) (Supplementary Fig. 8b).

We further compared the cellular fluorescence intensities of each FR/dye pair as measured using either a four-round (Fig. 3a) or a two-round (Fig. 3c) protocol. Even though each dye molecule was only incubated for 5 min following the rapid two-round approach, our results showed that the fluorescence signals of each FR/dye pair were still quite similar (∼73–94%) to those from the four-round imaging protocol with 20-min dye incubation time (Supplementary Fig. 8c). Meanwhile, minimal cytotoxicity was shown following both the four-round and two-round sequential imaging process (Fig. 3d). All these above experiments demonstrated that the seqFRIES system can indeed be used for the rapid multiplexed imaging in living bacterial cells.

### Multiplexed sequential imaging in mammalian cells

We next wanted to test if our seqFRIES system can also be applied for multiplexed imaging inside mammalian cells. To improve the cellular RNA expression level, we used a Tornado expression system^32^ to separately encode a circularized F30-Broccoli (abbreviated as cBroccoli), a circular F30-Corn (cCorn), a circular F30-DNB (cDNB), or a circularized F30-Pepper (cPepper) RNA within pAVU6+27-Tornado vectors. After transfecting the cBroccoli, cCorn, cDNB, or cPepper plasmid, respectively, into HEK293T cells, the DFHBI-1T, DFHO, TMR-DN, or HBC620 dye was separately added and incubated for 30 min before imaging. Similar to that shown in the bacterial cells, the fluorescence signals of these dye molecules can only be activated in HEK293T cells that express the corresponding fluorogenic RNA aptamers (Fig. 4a, b). The cBroccoli/DFHBI-1T, cCorn/DFHO, cDNB/TMR-DN, and cPepper/HBC620 FR/dye pairs can still be imaged orthogonally in mammalian cells.

**Fig. 4.**
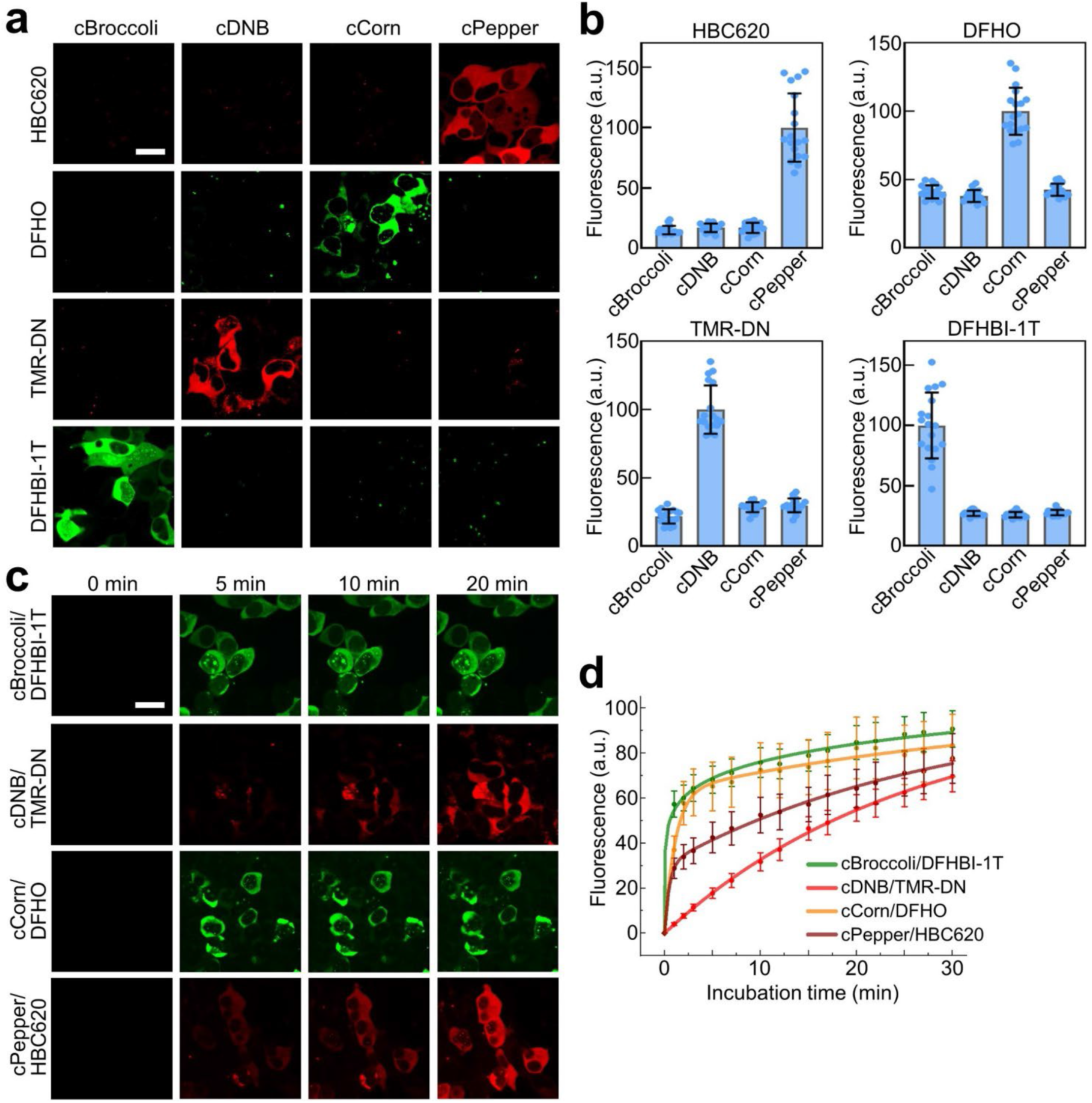
Orthogonality and fluorescence activation kinetics of FR/dye pairs in living mammalian cells. **a** Fluorescence images were taken in HEK293T cells that express circularized cBroccoli, cDNB, cCorn, or cPepper RNAs after a 30-min incubation with 40 μM DFHBI-1T, 1 μM TMR-DN, 40 μM DFHO, or 5 μM HBC620 dye molecules. Scale bar, 20 μm. **b** Cellular fluorescence intensities of the orthogonality experiments shown in panel (**a**) were also quantified in ∼20 individual cells in each case. Shown are the mean and SD values from cellular images taken from at least three independent replicates. **c** Cellular fluorescence signals in HEK293T cells that express circularized cBroccoli, cDNB, cCorn, or cPepper RNAs were monitored after adding 40 μM DFHBI-1T, 1 μM TMR-DN, 40 μM DFHO, or 5 μM HBC620 dye molecules at 0 min. Scale bar, 20 μm. **d** Cellular fluorescence activation kinetics as measured in ∼20 individual HEK293T cells in each case. Shown are the mean and the standard error of the mean (SEM) values from cellular images taken from at least three independent replicates.

We also investigated the fluorescence activation and deactivation kinetics of each FR/dye pair in HEK293T cells. As shown in Fig. 4c, d, immediately after adding the DFHBI-1T or DFHO dye, very fast fluorescence activation was observed in cells that express cBroccoli or cCorn RNAs, with half maximal fluorescence signals shown within ∼1 min. Similarly, upon the addition of HBC620, cPepper-expressing cells exhibited a rapid increase in cellular fluorescence intensities, in ∼9 min, half maximal fluorescence value could be observed. In the case of the cDNB/TMR-DN pair, similar to that shown in the *E. coli* cells, a gradual increase in cellular fluorescence was noticed, with ∼50% of the maximal signals detected at ∼18 min.

In the matter of fluorescence deactivation kinetics measurement, after pre-incubating FR-expressing HEK293T cells with the cognate dyes for 30 min, changes in the cellular fluorescence signals were monitored during five rounds of 1-min medium swap with dye-free DPBS, and the fluorescence images were also collected afterwards for another 5 min of incubation (Fig. 5a). Right after the five rounds of washing, cBroccoli/DFHBI-1T, cCorn/DFHO, and cDNB/TMR-DN signals were rapidly reduced respectively to ∼9%, 20%, and 23% of the original fluorescence intensities (Fig. 5b). While in contrast, cPepper/HBC620 fluorescence was more difficult to strip under this washing condition: ∼40% of the original signals remain in the cells. To solve this problem, we tried to apply the above-mentioned photobleaching-based stripping approach via a 2–4 min irradiation with a 1 mW/cm^2^ low power 561 nm laser. Indeed, cellular cPepper/HBC620 fluorescence signals could be rapidly reduced to ∼10% of original after this photo-stripping operation (Fig. 5c). Meanwhile, the minimal influence of this photobleaching step on the cell viability was validated via a SYTOX Blue dead cell staining assay (Supplementary Fig. 9a). As a result, fast fluorescence activation and deactivation of all these four FR/dye pairs can be achieved inside mammalian cells.

**Fig. 5.**
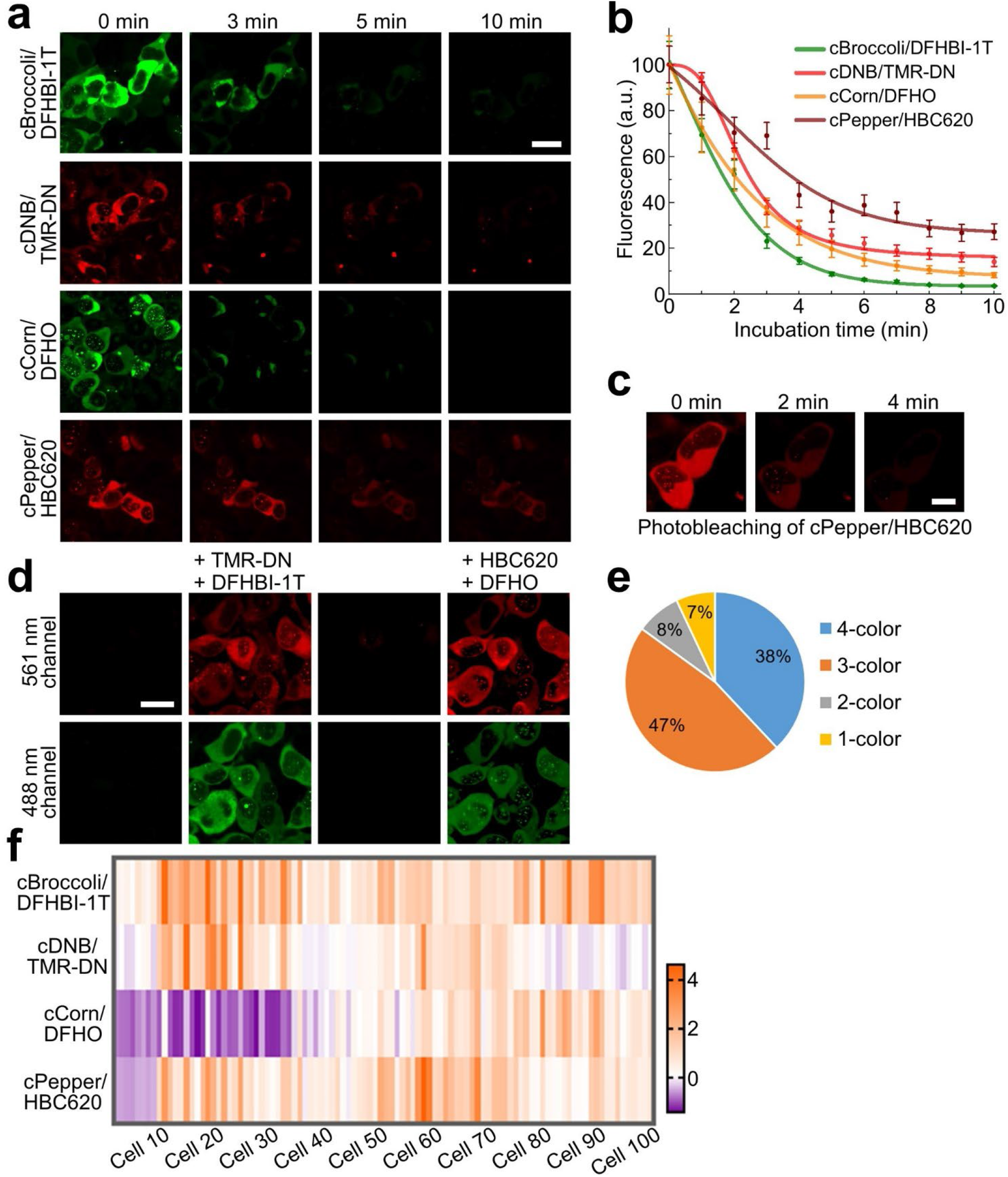
Fluorescence deactivation kinetics and sequential multiplexed imaging in HEK293T cells. **a** After incubating HEK293T cells that express cBroccoli, cDNB, cCorn, or cPepper for 30 min with 40 μM DFHBI-1T, 1 μM TMR-DN, 40 μM DFHO, or 5 μM HBC620 dye molecules, cellular fluorescence images were taken during and after five rounds of 1-min DPBS washing. Scale bar, 20 μm. **b** Cellular fluorescence deactivation kinetics as measured in ∼20 individual HEK293T cells in each case. Shown are the mean and SEM values from cellular images taken from at least three independent replicates. **c** Stripping of cellular fluorescence signals via brief photobleaching. After incubating HEK293T cells that express circularized cPepper RNAs for 30 min with 5 μM HBC620, cellular fluorescence images were taken after 2–4 min of irradiation with a 1 mW/cm^2^ power 561 nm laser. Scale bar, 10 μm. **d** Fluorescence images were taken in HEK293T cells that express all four FRs (cFDB-cFPC) after a sequential 20-min incubation with a mixture of 40 μM DFHBI-1T and 1 μM TMR-DN, or a mixture of 40 μM DFHO and 5 μM HBC620. Three times of 2-min DPBS washing were performed to strip cellular fluorescence. Scale bar, 15 μm. **e** Percentage of HEK293T cells that exhibit 1–4 channel activated fluorescence signals as measured in 100 cells after sequential imaging process shown in panel (**d**). Cellular images were taken from at least three independent replicates. **f** Heatmap of four FR/dye pair fluorescence signals from 100 individual HEK193T cells as measured via the seqFRIES process shown in panel (**d**). The values in the pseudo-color scale indicated the relative brightness of each imaging channel, calculated by subtracting the threshold fluorescence from the absolute cellular fluorescence values, and then divided by the standard deviation of the absolute fluorescence of each row. The positive values indicated the “on” fluorescence, while negative values meant the “off” fluorescence.

To test the efficiency of sequential multiplexed imaging, we further cloned both a circularized F30-DNB/Broccoli RNA (abbreviated as cFDB) and a circular F30-Pepper/Corn construct (cFPC) into an AIO-Puro-based plasmid. After transfecting the cFDB-cFPC plasmid into HEK293T cells, we first incubated the cells with DFHBI-1T and TMR-DN dyes for 20 min, and after imaging, three times of 2-min DPBS washing was performed to remove DFHBI-1T and TMR-DN before adding DFHO and HBC620 for another 20-min incubation for the second round of imaging (Fig. 5d). After this whole process of seqFRIES, indeed the majority (∼85%) of the transfected HEK293T cells exhibited fluorescence signals from at least three or four FR/dye pairs (Fig. 5e). While another ∼7% and 8% of cells displayed only one or two colors. In those HEK293T cells that exhibit at least one set of FR/dye fluorescence, the Broccoli/DFHBI-1T signals could be observed in near 100% of cells, while in contrast, the Corn/DFHO signals were more difficult to be visualized, which was activated in only ∼55% of the HEK293T cells (Fig. 5f). In agreement with these results, relatively poor cellular fluorescence correlations (Pearson’s r< 0.27) were observed between cCorn/DFHO and the other three FR/dye channels. In comparison, nice positive correlations among cPepper/HBC620, cDNB/TMR-DN, and cBroccoli/DFHBI-1T were shown in these same cells with a Pearson’s r value in the range of 0.48– 0.71 (Supplementary Fig. 9b). Still, these results indicated the successful performance of seqFRIES for multi-color sequential imaging in living mammalian cells.

### seqFRIES-mediated cellular detection of different target analytes

After demonstrating the cellular functions of the seqFRIES system, we lastly wanted to test if this sequential reporting system can be used for detecting different target analytes. For this purpose, two bacterial second messengers, cyclic diguanylate (c-di-GMP) and guanosine tetraphosphate (ppGpp), were chosen as the targets because of their critical regulatory functions in bacterial physiology and stress responses^33–35^. In addition, we have engineered a Pepper-based sensor for detecting S-adenosyl-L-methionine (SAM) (Supplementary Fig. 10a), a cellular methyl donor that can be converted to S-adenosyl-L-homocysteine (SAH) during cellular methylation process. Considering the importance of SAM:SAH ratio as an indicator of cellular methylation potential^36,37^, we have decided to study the possibility of developing orthogonal FR-based biosensors for detecting c-di-GMP, ppGpp, SAM, and SAH.

Our sensor system contains a Broccoli-based ppGpp sensor (named as **B-P4G**)^38^, a DNB-based c-di-GMP sensor (**D-CDG**)^29^, a Pepper-based SAM sensor (**P-SAM**), and a Corn-based SAH sensor (**C-SAH**)^39^. The performance of each FR-based biosensor was first tested in a mixture together with another three sensors. As shown in Fig. 6a, b, both the target binding affinity and selectivity of all these FR-based sensors were well maintained in the mixture. Physiological concentrations of c-di-GMP, ppGpp, SAM, and SAH can be potentially detected based on the dose-response curves of these sensors. The four orthogonal FR/dye pairs used for developing seqFRIES are thus also potentially capable of being engineered into biosensors, which can function independently in the same solution.

**Fig. 6.**
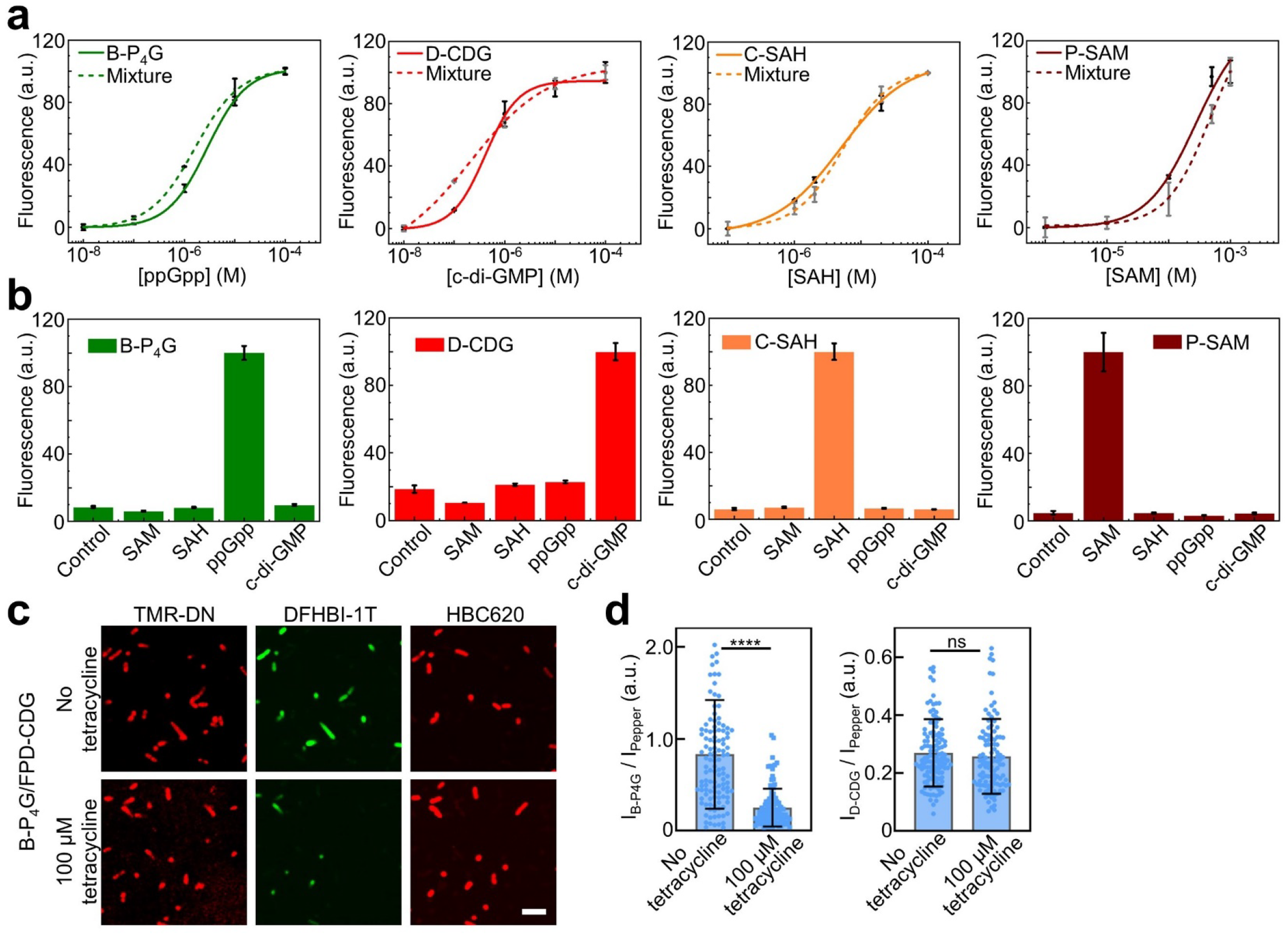
seqFRIES-based metabolite and signaling molecule sensors. **a** Dose-response curves for the fluorescence detection of ppGpp, c-di-GMP, SAM, and SAH by the optimal B-P4G, D-CDG, P-SAM, and C-SAH sensors, either using each corresponding sensor, or a mixture of all four sensors at 1–5 μM concentration. Shown are the mean and SEM values from at least three independent replicates. **b** Selectivity of each FR-based sensors as measured in a solution containing 1–5 μM RNA, 20 μM DFHBI-1T, 0.5 μM TMR-DN, 2 μM DFHO, or 2 μM HBC620, and 100 μM indicated compounds for B-P4G, D-CDG, P-SAM, or 10 μM indicated compounds for C-SAH. Shown are the mean and SEM values from at least three independent replicates. **c** Fluorescence images were taken in BL21 Star (DE3) cells that co-express B-P4G and FPD-CDG RNAs for detecting both ppGpp and c-di-GMP signaling molecules. 200 μM DFHBI-1T and 1 μM TMR-DN were added together and incubated for 30 min before the first round of imaging, after three times of 5-min washing, the second round of imaging was performed after a 30-min incubation with 1 μM HBC620. The same seqFRIES procedure was performed in the presence or absence of 100 μM tetracycline for studying antibiotic-induced changes in cellular ppGpp and c-di-GMP levels. Scale bar, 5 μm. **d** Using cellular Pepper/HBC620 signals as the reference, the fluorescence ratios of B-P4G/DFHBI-1T to Pepper/HBC620 (I_B-P4G_/I_Pepper_) and D-CDG/TMR-DN to Pepper/HBC620 (I_D-CDG_/I_Pepper_) were measured in ∼100 individual cells with and without the presence of 100 μM tetracycline. Shown are the mean and SD values from cellular images taken from at least three independent replicates. Two-tailed student’s t-test: ****, p<0.0001; ns, not significant, p>0.05.

To further study the cellular performance of these RNA-based sensors for sequential multiplexed detection, we encoded F30-scaffolded D-CDG and Pepper (**FPD-CDG**) into one T7 expression cassette, and B-P4G into another T7 expression cassette within the same pETDuet plasmid (Supplementary Fig. 10b). In this system, Pepper fluorescence was introduced as a reference signal to normalize RNA expression level variations in individual *E. coli* cells, while D-CDG and B-P4G were used to respectively detect cellular c-di-GMP and ppGpp levels. The cellular performance of D-CDG and B-P4G sensors have been separately validated in our previous studies^29,38^. Our goal here is to enable the first direct live-cell measurement of both c-di-GMP and ppGpp, two critical signaling molecule targets that have been shown to collaboratively facilitate bacterial survival under stress conditions, such as the antibiotic treatment^40–44^.

After co-expressing FPD-CDG and B-P4G inside BL21 Star (DE3) cells, we first incubated the cells with DFHBI-1T and TMR-DN dyes for 30 min to simultaneously detect cellular c-di-GMP and ppGpp levels. Afterwards, three times of 5-min washing was used to strip cellular fluorescence before the next round of imaging for Pepper/HBC620 reference signals. Indeed, bright cellular D-CDG/TMR-DN, B-P4G/DFHBI-1T, and Pepper/HBC620 fluorescence could be observed during this sequential imaging-and-stripping process (Fig. 6c). The same seqFRIES procedure was also performed in the presence of 100 μM tetracycline for studying the effect of antibiotic treatment on the cellular ppGpp and c-di-GMP concentrations. By normalizing RNA expression levels in each individual cell via the Pepper/HBC620 signals, the ratiometric fluorescence intensity of I_B-P4G_/I_Pepper_ and I_D-CDG_/I_Pepper_ was used here to respectively detect the cellular changes in ppGpp and c-di-GMP levels after the tetracycline antibiotic treatment. An ∼70% lower I_B-P4G_/I_Pepper_ signals were shown upon adding 100 μM tetracycline, while cellular c-di-GMP levels were not significantly affected in the same test (Fig. 6d). To rule out the possibility that the addition of tetracycline may influence the cellular fluorescence signals of Broccoli/DFHBI-1T and DNB/TMR-DN, a control experiment was performed using FDB-FPC-expressing cells. Following a same seqFRIES procedure, none of the Broccoli/DFHBI-1T, DNB/TMR-DN, and Pepper/HBC620 fluorescence signals were affected by the addition of 100 μM tetracycline (Supplementary Fig. 10c, d). All these results suggest that these *E. coli* cells tend to reduce the accumulation of intracellular ppGpp levels to survive under antibiotic stress, while c-di-GMP may not play a direct role in this process. We expect that some more detailed mechanism studies will be followed in the future.

## Discussions

We herein introduced a seqFRIES platform for live-cell sequential multiplexed imaging. The seqFRIES system functions by using multiple rounds of imaging-and-stripping steps to overcome the spectral overlap issues in traditional fluorescent techniques. One unique feature of our fluorogenic RNA-based system is that the dye molecules can be reversibly separated from the genetically encoded RNA molecules, and as a result, the chemical dyes can be added externally and washed away easily afterward. In this study, we have identified several pairs of orthogonal fluorogenic RNA/dye conjugates, such as Broccoli/DFHBI-1T, Corn/DFHO, DNB/TMR-DN, and Pepper/HBC620, and showed that by adding only one or two dyes at a time, all these fluorescent pairs can be imaged inside the same individual cells without spectrally influencing each other.

We have also demonstrated that this novel multiplexed imaging strategy can function in both living bacterial and mammalian cells. The seqFRIES technique does not require cell fixation or permeabilization. The dye molecules are permeable to the living cells membranes and exhibit fast fluorescence activation and deactivation kinetics with their cognate RNA aptamers inside the cells. The optimized high efficiency labeling-and-stripping procedure allowed us to image four fluorogenic RNA/dye pairs in cells in just 20 minutes. Our results also demonstrated that the seqFRIES system can enable single-cell analysis of different target analytes, via repeated fluorescence imaging in the same individual cells. Such an ability of performing single-cell analysis can potentially provide valuable information on the cell-to-cell variations in their molecular signatures, which are usually masked in bulk analysis.

Genetically encoded fluorogenic RNA-based biosensors have become popular tools for the detection and monitoring of cellular targets. Compared to small molecule- or fluorescent protein-based sensors, the unique advantages of these fluorogenic RNA-based sensing platforms are their high programmability and generalizability. These fluorogenic RNAs can function as imaging tags or via sequence-specific hybridizations for the cellular tracking of various mRNA and non-coding RNA analytes^4,5,19^. RNA aptamers can also be selected for a wide range of cellular targets, including different proteins, metabolites, ions, and signaling molecules^45,46^. These specific aptamers can be further modularly engineered into targeted fluorescent sensors. The versatility of this RNA-based seqFRIES system will make it easily applied for the future detection of a large number of target cellular analytes, at the single-cell level.

To potentially allow this seqFRIES system to detect more than four targets, the identification of additional orthogonal fluorogenic RNA/dye pairs are still required to expand the current multiplexing platform. Indeed, many fluorogenic RNA/dye pairs have been discovered recently, such as SiRA/SiR and o-Coral/Gemini-561^47,48^, which can be potentially orthogonal to our current FR/dye pairs in this project. Furthermore, by using an internal reference FR/dye pair, e.g., Pepper/HBC620, to normalize cell-to-cell variations in RNA expression levels, we have partially validated the possibility of applying the seqFRIES system in quantitative cellular analysis. With further optimization and expansion of the orthogonal FR/dye pairs, we expect that the seqFRIES platform can be broadly applied for multiplexed live-cell imaging and quantitative molecular profiling to reveal molecular correlations and provide a better understanding of mysterious cellular processes.

## Methods

### Reagents

All the chemicals were purchased from MilliporeSigma or Fisher Scientific unless indicated otherwise. Commercial reagents were directly used without additional purification. HBC620, 4-(2-hydroxyethyl-methylamino)-benzylidene-cyanophenylacetonitrile analog, was purchased from GlpBio (Montclair, CA). YO3-biotin, an oxazole yellow derivative, was purchased from Applied Biological Materials Inc (Vancouver, Canada). S-(5’-Adenosyl)-L-methionine chloride was purchased from Cayman Chemical (Ann Arbor, MI). 3,5-difluoro-4-hydroxybenzylidene imidazolinone-2-oxime (DFHO) was a gift from Samie R. Jaffrey’s lab at Weill Cornell Medicine.

### Synthesis of oligonucleotides

DNA oligonucleotides were synthesized and cartridge-purified by Integrated DNA Technologies (Coralville, IA) or W. M. Keck Oligonucleotide Synthesis Facility (Yale University School of Medicine). Oligonucleotides were then dissolved at 100 μM concentration in TE buffer (10 mM Tris-HCl, 0.1 mM EDTA, pH= 7.5) and stored at -20°C. Double-stranded DNAs were made by PCR amplification using an Eppendorf Mastercycler. The PCR products were cleaned using a QIAquick PCR purification kit (Qiagen, Germantown, MD). Concentration measurement of nucleic acids was performed using a NanoDrop One UV-vis spectrophotometer. RNA synthesis was conducted via *in vitro* transcription using a HiScribe™ T7 high yield RNA synthesis kit (New England BioLabs, NEB, Ipswich, MA). After DNase I (RNase-free) (NEB) treatment to remove the DNA strands, the transcribed RNAs were column-purified and further verified by running 10% denaturing polyacrylamide gel electrophoresis. All the synthesized RNA products were separated into aliquots and stored at -20°C for immediate usage or at -80°C for long-time storage. NUPACK and mFold online software were used to simulate and design all the RNA structures.

### *In vitro* fluorescence assay

All the *in vitro* fluorescence measurements were performed with a PTI fluorimeter (Horiba, New Jersey, NJ) at room temperature (23°C). Fluorescence signals were measured in a buffer consisting of 40 mM Tris, 5 mM MgCl2, 100 mM KCl at pH= 7.6. The fluorescence emission spectra of Chili, Broccoli, Corn, DNB, Pepper, and MangoII were collected by exciting at 415 nm, 480 nm, 505 nm, 555 nm, 577 nm, and 580 nm, respectively. An Origin software was used to plot all these *in vitro* fluorescence data.

### Vector construction for bacterial imaging

Pepper, DNB, Corn, and FDB were cloned into a pETDuet-1 vector (EMD Millipore, Burlington, MA), respectively. This vector was first double digested with SgrAI and NdeI restriction enzymes (NEB) and purified from 1% agarose gel using a QIAquick Gel Extraction Kit (Qiagen, Germantown, MD). A double-stranded insert was digested at the same restriction sites, followed by ligation with the digested vector, using T4 DNA ligase (NEB). Chili and FPC were cloned respectively into the pETDuet-1 by double digestion and ligation following the same procedure described above, and the restriction enzymes used were NdeI and PacI (NEB). The pETDuet-Broccoli plasmid was a gift from Samie R. Jaffrey’s lab at Weill Cornell Medicine. F30-D-CDG/Pepper was cloned into the pETDuet plasmid via double digestion by SgrAI and SacII, and B-P4G was cloned into the same vector via NdeI and XhoI digestion. All the ligated products were then transformed into TOP10 chemical competent cells (Invitrogen) and screened based on ampicillin resistance. All the plasmids were extracted using a GeneJET Plasmid Miniprep Kit (Thermo Fisher Scientific) and confirmed via Sanger sequencing performed by Eurofins Genomics.

### Vector construction for mammalian imaging

For mammalian cell imaging, cFDB and cFPC were cloned into an AIO-Puro vector (Addgene). The DNA templates of cFDB and cFPC were respectively digested with BsaI and Bbs1 restriction enzymes (NEB) and ligated with the vector digested at the same restriction sites. cBroccoli, cDNB, cPepper, and cCorn were cloned into an pAVU6+27-Tornado vector (Addgene). The DNA templates of cBroccoli, cDNB, cPepper, and cCorn were double digested by NotI-HF and SacII (NEB), and ligated with the vector digested at the same restriction sites. All the ligated products were then transformed into TOP10 chemical competent cells (Invitrogen) and screened based on ampicillin resistance. All the plasmids were extracted using a GeneJET Plasmid Miniprep kit (Thermo Fisher Scientific) and confirmed by Sanger sequencing by Eurofins Genomics.

### Bacterial cell imaging and data analysis

Confocal imaging of *E. coli* cells was conducted according to a previously reported protocol^49^. The BL21 Star (DE3) cells were grown to OD600 of 0.4–0.5 at 37°C in LB media, and then 1 mM isopropyl β-D-1-thiogalactopyranoside (IPTG) was added for 4 h to induce cellular RNA transcription. All the fluorescence images were collected with a NIS-Elements AR software using a Yokogawa spinning disk confocal on a Nikon Eclipse-TI inverted microscope. Broccoli/DFHBI-1T and Corn/DFHO were excited with a 488 nm laser, and the emission was collected in the range of 500–550 nm via a filter set. Pepper/HBC620 and DNB/TMR-DN were imaged by a 561 nm laser, and the fluorescence emission was collected in the range of 575–625 nm. Chili was excited with a 405 nm laser, and the emission was collected in the range of 500–550 nm. All these images were obtained through a 60x or 100× oil immersion objective. Data analysis was performed with ImageJ software. Bacterial cells were identified by a MicrobeJ plugin using the local default setting. Only the fluorescence signals from surface well-attached rod-shaped bacterial cells were used for quantitative comparison. Data calculation and fitting were carried out using the Origin software.

### Mammalian cell culture and transfection

HEK293T/17 cells were purchased from ATCC and cultured in DMEM (high glucose) supplemented with 10% FBS and 1% penicillin-streptomycin under 5% CO2 at 37°C. The cell line was mycoplasma-free and split using 0.25% trypsin at ∼80% confluence. Transfection was achieved using the FuGENE HD transfection reagent (Promega) according to the manufacturer’s instructions. More specifically, 1 μg DNA plasmids and 3 μL FuGENE reagent were first mixed with 50 μL Gibco Opti-MEM medium and incubated at room temperature for 15 min. Then, 12 μL of the mixture was added to a chamber of poly-D-lysine-coated 8-chamber glass bottom slide (Cellvis, CA) for imaging. The same batch of HEK293T/17 cells without transfection were used as the negative control and added to another chamber of the same imaging slide. After 24 hours of transfection, these cells were ready for imaging.

### Mammalian cell imaging

HEK293T/17 cell imaging was also conducted using a Yokogawa spinning disk confocal on a Nikon Eclipse-TI inverted microscope. All the images were obtained through a 40× oil immersion objective. cBroccoli/DFHBI-1T and cCorn/DFHO were excited with a 488 nm laser, and the emission was collected in the range of 500–550 nm via a filter set. cPepper/HBC620 and cDNB/TMR-DN were imaged by a 561 nm laser, and the fluorescence emission was collected in the range of 575-625 nm. Data analysis was performed with ImageJ software. Cells were identified manually for quantitative analysis. Data calculation and fitting were carried out using the Origin software.

### Determination of fluorescence threshold values in HEK293T cells

Cellular fluorescence signals of untransfected HEK293T cells after a 30-min incubation with the corresponding dye molecules at a concentration of 40 μM DFHBI-1T, 1 μM TMR-DN, 40 μM DFHO, or 5 μM HBC620 are used to determine the background cellular fluorescence intensities for each imaging channel. The fluorescence threshold values of HEK293T cells were then calculated as the average background fluorescence (μ) from 10 untransfected cells plus two times of the standard deviation (σ) values of these background signals, i.e., “μ+2σ”. In each imaging channel, HEK293T cells that exhibit greater than μ+2σ cellular fluorescence intensities were considered as having this imaging channel “on”. Otherwise, the corresponding imaging channel was considered as “off”.

### Cytotoxicity measurement

Cytotoxicity measurement was performed by incubating BL21 Star (DE3) or HEK293T cells with 1 μM SYTOX Blue nucleic acid stain (Invitrogen) for 5 min before imaging by a 405 nm laser irradiation.

### Synthesis of tetramethylrhodamine-PEG3-dinitroaniline (TMR-DN)

The tetramethylrhodamine-PEG3-dinitroaniline (TMR-DN) was synthesized following a previously reported procedure^24^. To briefly describe, dinitroaniline-PEG3-amine (DN-PEG3-amine) was first synthesized by adding a 10 mL dichloromethane (DCM) solution of dinitrofluorobenzene (1.0 g, 5.4 mmol) dropwise into a solution of 2,2’-(ethylenedioxy)bisethylenediamine (4.67 g, 32.4 mmol) in 20 mL of DCM at 0°C. The reaction mixture was then stirred at room temperature for 30 min. Afterwards, washed the reaction mixture with 100 mL water, and recovered DN-PEG3-amine in the DCM phase, which was then extracted to the aqueous phase in the presence of 0.1 M HCl. After adjusting the pH of the aqueous solution to ∼12–13 and mixing it with DCM, the DN-PEG3-amine product was extracted back to the DCM phase and dried overnight. The product was used without further purification for the next step of reaction. HR-ESI (positive): calculated 315.1299, found 315.1594 for C12H19N4O6.

To synthesize TMR-DN, the above-prepared DN-PEG3-amine (1.8 mg, 5.7 μmol) was dissolved in 50 μL of dimethylformamide (DMF) and added dropwise to a solution of 5-carboxy-tetramethylrhodamine-N-hydroxysuccinimide (1.0 mg, 1.9 μmol) in 100 μL of DMF. The reaction mixture was then stirred at room temperature for 15 min. The reaction mixture was purified on a reversed-phase liquid chromatograph with a C18 column and a mobile phase composition of 60% acetonitrile and 0.1% trifluoroacetic acid to yield TMR-DN (0.53 mg, 38% yield). HR-ESI (positive): calculated 727.2722, found 727.3671 for C37H39N6O10.

## Data availability

The data supporting the findings of this study are available upon request to the corresponding author.

## Acknowledgements

The authors gratefully acknowledge the support from Chan Zuckerberg Initiative Dynamic Imaging program, NSF CAREER award #1846152, Alfred P. Sloan Research Fellowship, Camille Dreyfus Teacher-Scholar Award and UMass Amherst start-up grant to M.Y. R.Z. was also supported by an NIH T32GM139789 Traineeship. Z.S. and Y.B. were also supported by a Paul Hatheway Terry Scholarship. We are grateful to Dr. James Chambers for the assistance in fluorescence imaging. The authors also thank other members of the You Lab for useful discussion and valuable comments.

## Author contributions

M.Y., R.W., and K.R. conceived the study. R.Z. performed all the in vitro and bacterial cell experiments with help from R.W., Z.S., R.P., and S.S. Y.L. and R.Z. performed mammalian cell measurements, with help from L.M. and Q.T. Y.B., Z.S., and R.Z. synthesized the TMR-DN dye. R.Z. and R.W. analyzed the data with input from K.R., Z.X., and Q.T. M.Y. supervised all aspects of the research. All authors were involved in the discussion of the data. R.Z. and M.Y. wrote the manuscript with input from all the authors.

## Additional information

### Supplementary information

The online version contains supplementary material available.

