## Supplementary Information for "Multiplexed Sequential Imaging in Living Cells with Orthogonal Fluorogenic RNA Aptamer/Dye Pairs"

[†] These authors contribute equally

### Contents

#### Supplementary Figures

**Fig. S1**. *In vitro* identification of orthogonal fluorogenic RNA (FR)/dye pairs.
**Fig. S2**. Effect of RNA aptamer mixture and concentration on the fluorescence signals.
**Fig. S3**. Fluorescence activation kinetics of FR/dye pairs in live *E. coli* cells.
**Fig. S4**. Fluorescence deactivation kinetics of FR/dye pairs in live *E. coli* cells.
**Fig. S5**. Stripping of cellular fluorescence signals via photobleaching or competitive binding.
**Fig. S6**. Co-expression of four FRs in a pETDuet vector.
**Fig. S7**. Sequential imaging feasibility of FR/dye pairs in live *E. coli* cells.
**Fig. S8**. Simultaneous detection of two FR/dye pairs to shorten the overall imaging time.
**Fig. S9**. Measurement of cytotoxicity and multiplexed imaging correlations in HEK293T cells.
**Fig. S10**. seqFRIES-mediated metabolite and signaling molecule sensors.

#### Supplementary Table

**Table S1**. List of RNA sequences used in this work.

### Supplementary Figures

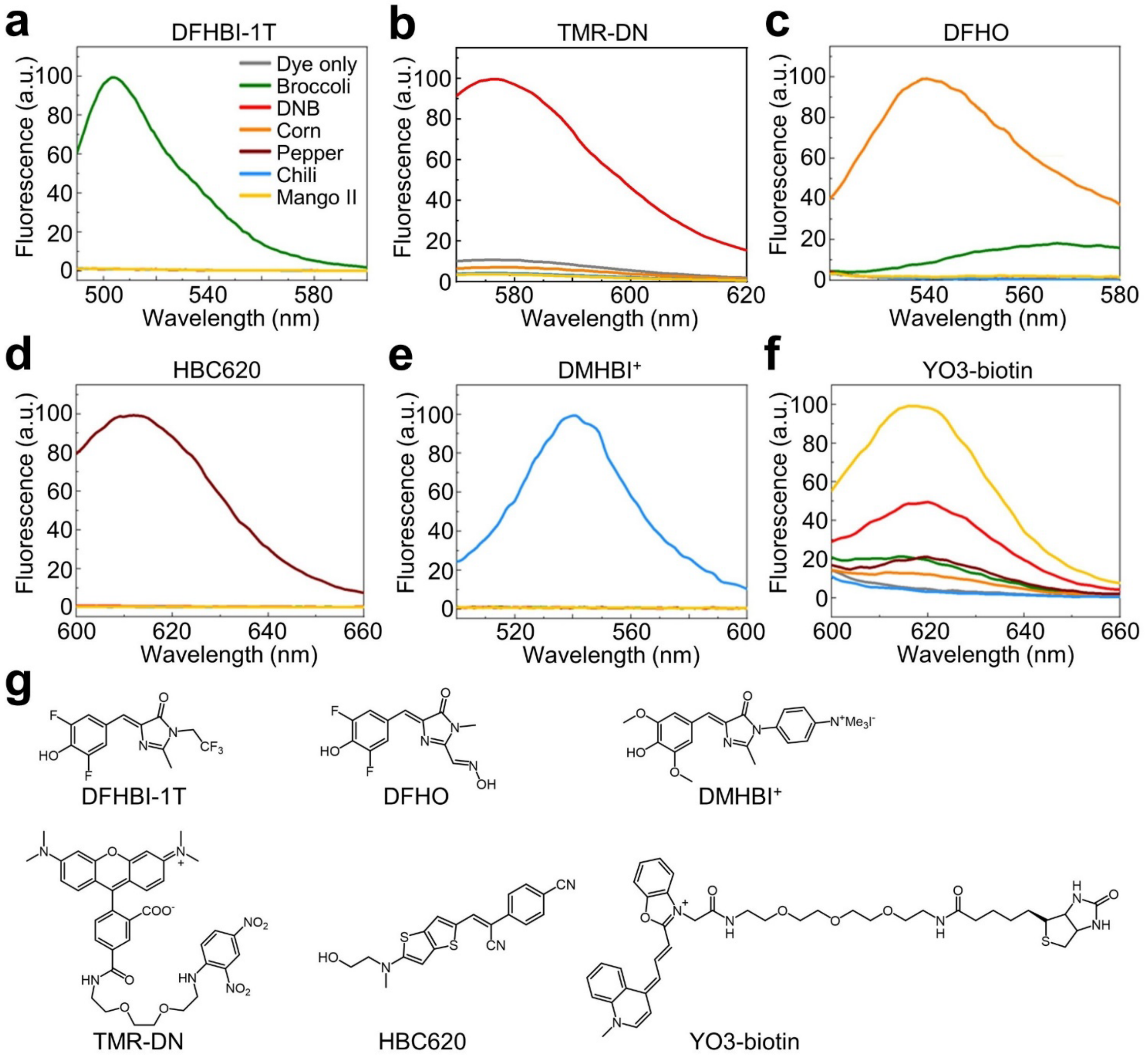


**Fig. S1.** ***In vitro*** **identification of orthogonal fluorogenic RNA (FR)/dye pairs.** (**a**) Fluorescence spectra as measured upon 480 nm excitation of a solution containing 20 μM DFHBI-1T and 1 μM corresponding FR. (**b**) Fluorescence spectra as measured upon 555 nm excitation of a solution containing 0.5 μM TMR-DN and 2.5 μM corresponding FR. (**c**) Fluorescence spectra as measured upon 505 nm excitation of a solution containing 2 μM DFHO and 5 μM corresponding FR. (**d**) Fluorescence spectra as measured upon 577 nm excitation of a solution containing 1 μM HBC620 and 1 μM corresponding FR. (**e**) Fluorescence spectra as measured upon 413 nm excitation of a solution containing 1 μM DMHBI$^+$ and 1 μM corresponding FR. (**f**) Fluorescence spectra as measured upon 580 nm excitation of a solution containing 100 nM YO3-biotin and 1 μM corresponding FR. Shown are the representative results from at least three independent replicates. (**g**) The chemical structure of each dye molecule used in this study.

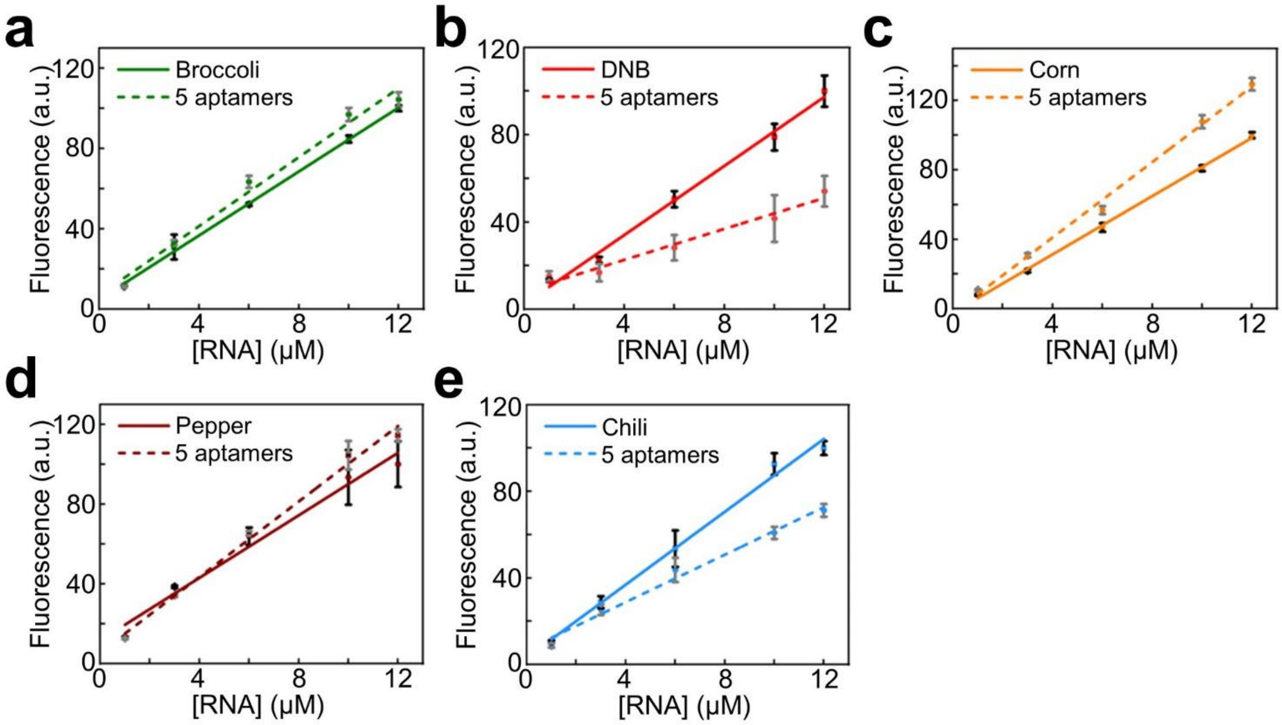


**Fig. S2. Effect of RNA aptamer mixture and concentration on the fluorescence signals.** "5 aptamers" indicates a mixture of five FRs at the same concentration, including Broccoli, DNB, Corn, Pepper, and Chili. (**a**) RNA concentration-dependent fluorescence signals ($\lambda_{ex}/\lambda_{em}$, 480/503 nm) as measured in a solution containing 20 µM DFHBI-1T. (**b**) RNA concentration-dependent fluorescence signals ($\lambda_{ex}/\lambda_{em}$, 555/582 nm) as measured in a solution containing 1 µM TMR-DN. (**c**) RNA concentration-dependent fluorescence signals ($\lambda_{ex}/\lambda_{em}$, 505/545 nm) as measured in a solution containing 12 µM DFHO. (**d**) RNA concentration-dependent fluorescence signals ($\lambda_{ex}/\lambda_{em}$, 577/612 nm) as measured in a solution containing 12 µM HBC620. (**e**) RNA concentration-dependent fluorescence signals ($\lambda_{ex}/\lambda_{em}$, 415/540 nm) as measured in a solution containing 12 µM DMHBI$^{+}$. Shown are the mean and standard deviation (SD) values from at least three independent replicates.

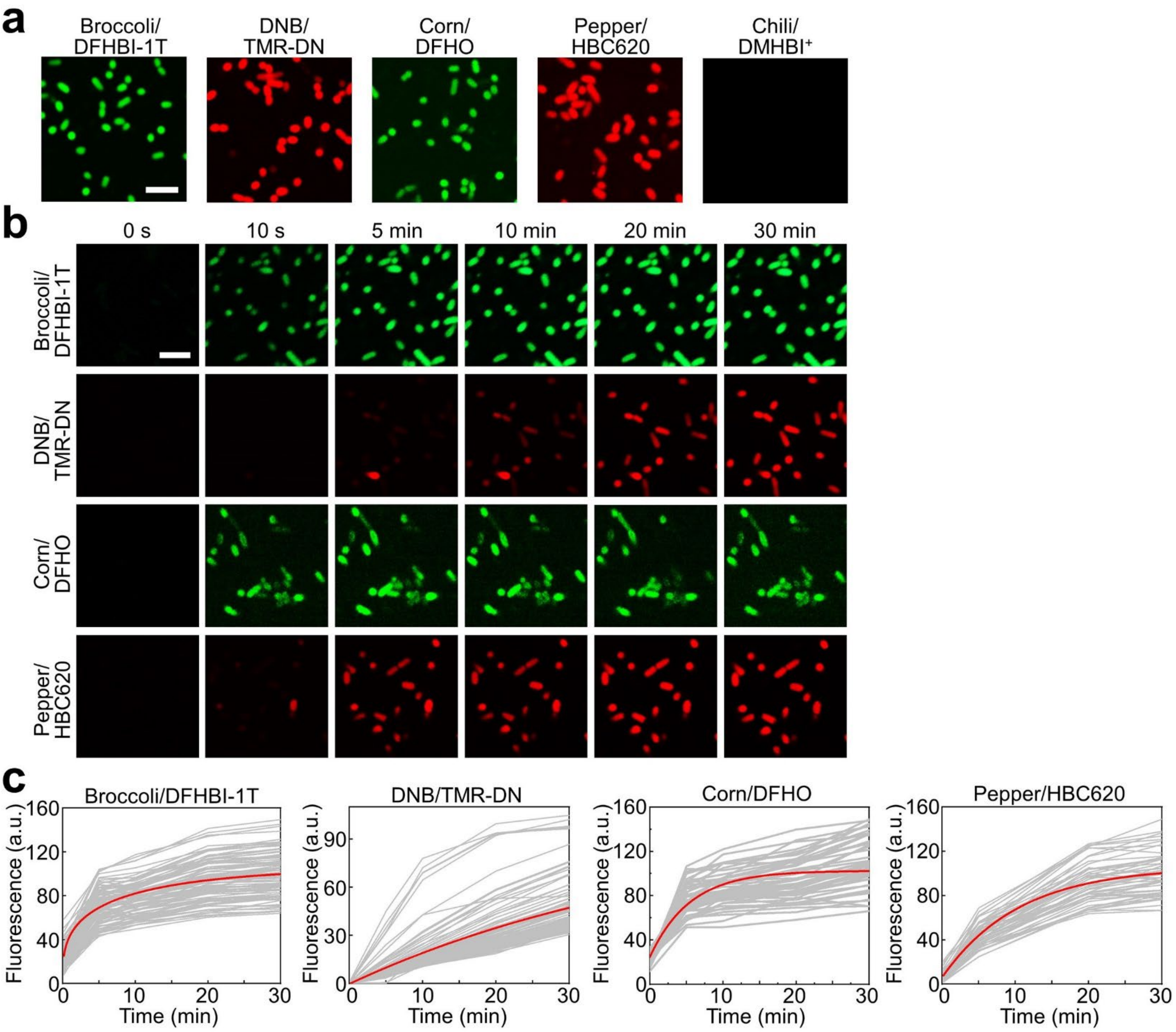


**Fig. S3. Fluorescence activation kinetics of FR/dye pairs in live *E. coli* cells.** (**a**) Fluorescence images were taken in BL21 Star (DE3) cells that express Broccoli, DNB, Corn, Pepper, or Chili RNAs after a 30-min incubation with 200 μM DFHBI-1T, 1 μM TMR-DN, 10 μM DFHO, 1 μM HBC620, or 5 μM DMHBI$^+$ dye molecules. Scale bar, 5 μm. (**b**) Cellular fluorescence signals were monitored for 30 min after adding 200 μM DFHBI-1T, 1 μM TMR-DN, 10 μM DFHO, or 1 μM HBC620 dye molecules into BL21 Star (DE3) cells expressing Broccoli, DNB, Corn, or Pepper RNAs. Scale bar: 5 μm. (**c**) Corresponding cellular fluorescence signals were plotted by measuring ~50–100 individual cells from three independent replicates. Each gray line represents the fluorescence signals imaged over time within an individual cell. Averaged cellular fluorescence signals were shown and fitted as the red line, which was also used to estimate the equilibrium cellular fluorescence intensities (set as 100).

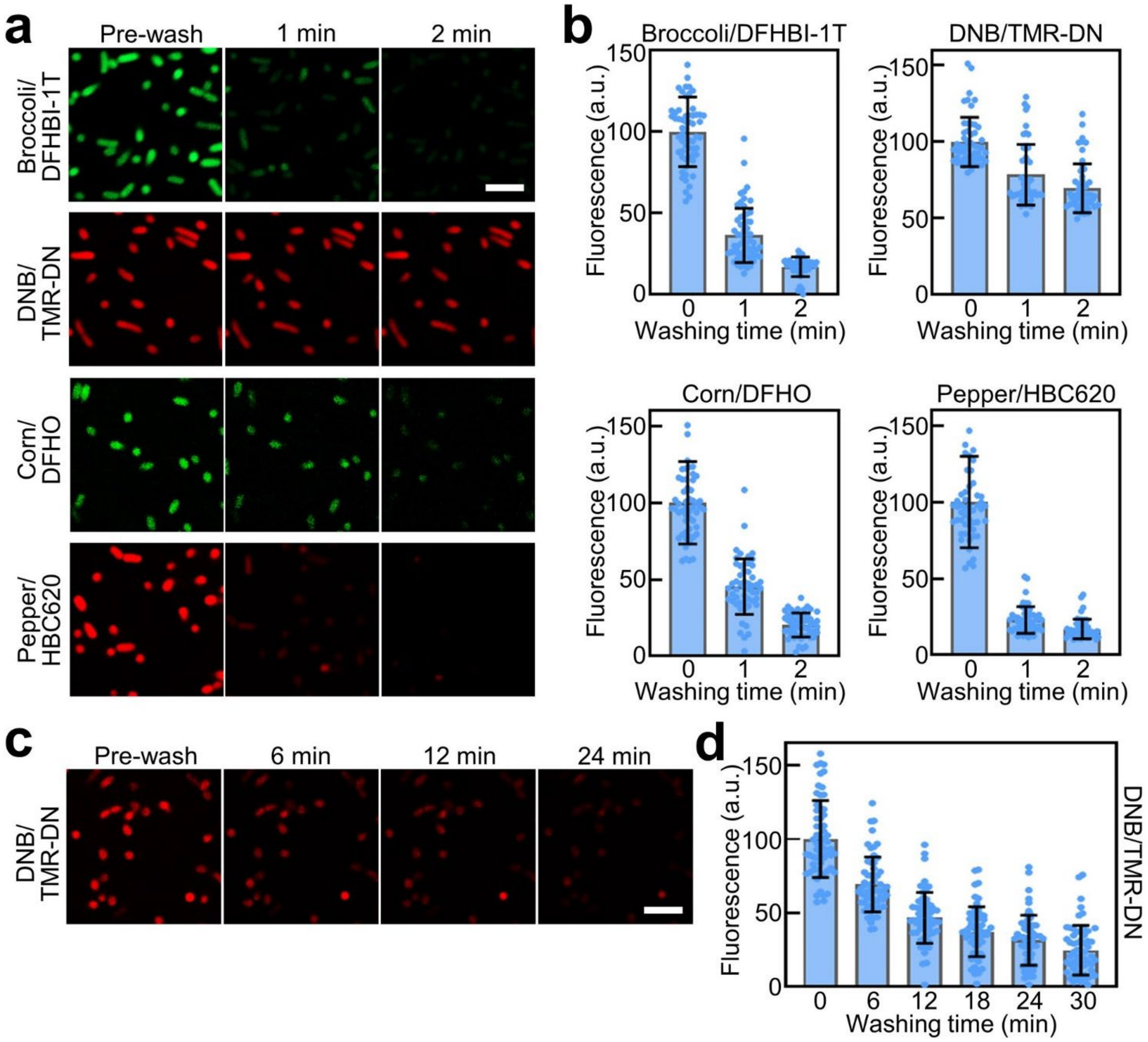


**Fig. S4. Fluorescence deactivation kinetics of FR/dye pairs in live *E. coli* cells.** (**a**) After incubating the BL21 Star (DE3) cells that express Broccoli, DNB, Corn, or Pepper RNAs for 50 min with 200 μM DFHBI-1T, 1 μM TMR-DN, 10 μM DFHO, or 1 μM HBC620 dye molecules, cellular fluorescence images were taken after once or twice of 1-min washing using a dye-free Dulbecco's phosphate-buffered saline (DPBS) buffer. Scale bar, 5 μm. (**b**) Cellular fluorescence intensities as measured in ~50 individual BL21 Star (DE3) cells before and after once or twice of 1-min washing with DPBS. Shown are the mean and SD values from cellular images taken from at least three independent replicates. (**c**) After incubating DNB-expressing BL21 Star (DE3) cells for 50 min with 1 μM TMR-DN, cellular fluorescence images were taken after 1–5 times of 6-min washing with fresh DPBS buffer. Scale bar, 5 μm. (**d**) Cellular fluorescence intensities as measured in ~50 individual DNB-expressing BL21 Star (DE3) cells before and after 1–5 times of 6-min washing. Shown are the mean and SD values from cellular images taken from at least three independent replicates.

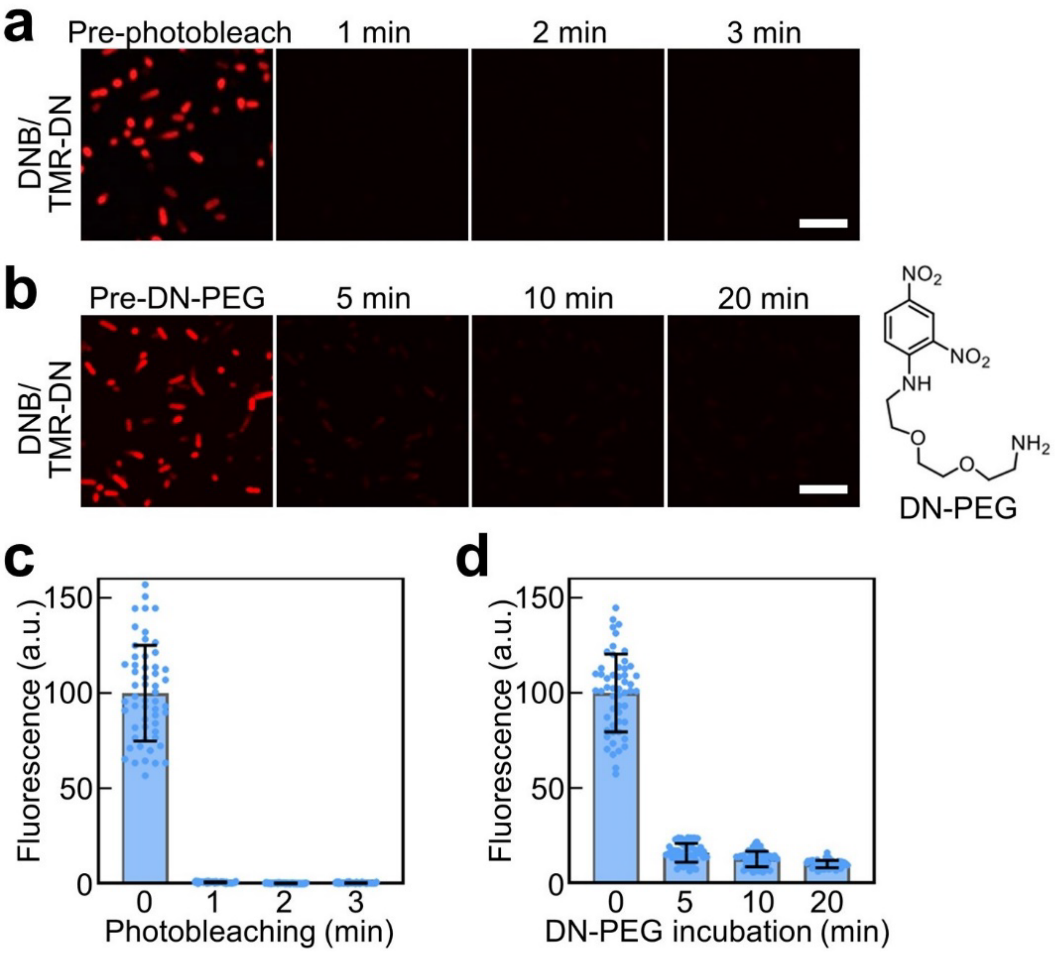


**Fig. S5. Stripping of cellular fluorescence signals via photobleaching or competitive binding.** (**a**) After incubating DNB-expressing BL21 Star (DE3) cells for 50 min with 1 μM TMR-DN, cellular fluorescence images were taken after 1–3 min of irradiation with a 1 mW/cm$^2$ power 561 nm laser. Scale bar, 5 μm. (**b**) After incubating DNB-expressing BL21 Star (DE3) cells for 50 min with 1 μM TMR-DN, cellular fluorescence images were taken after 5–20 min incubation with 20 μM nonfluorescent DN-PEG ligand to compete with TMR-DN for their binding with the DNB aptamer. The chemical structure of DN-PEG was also shown. Scale bar, 5 μm. (**c**) Cellular fluorescence intensities as measured in ~50 individual DNB-expressing BL21 Star (DE3) cells before and after 1–3 min of 561 nm laser irradiation. (**d**) Cellular fluorescence intensities as measured in ~50 individual DNB-expressing BL21 Star (DE3) cells before and after 5–20 min incubation with 20 μM DN-PEG. Shown are the mean and SD values from cellular images taken from at least three independent replicates.

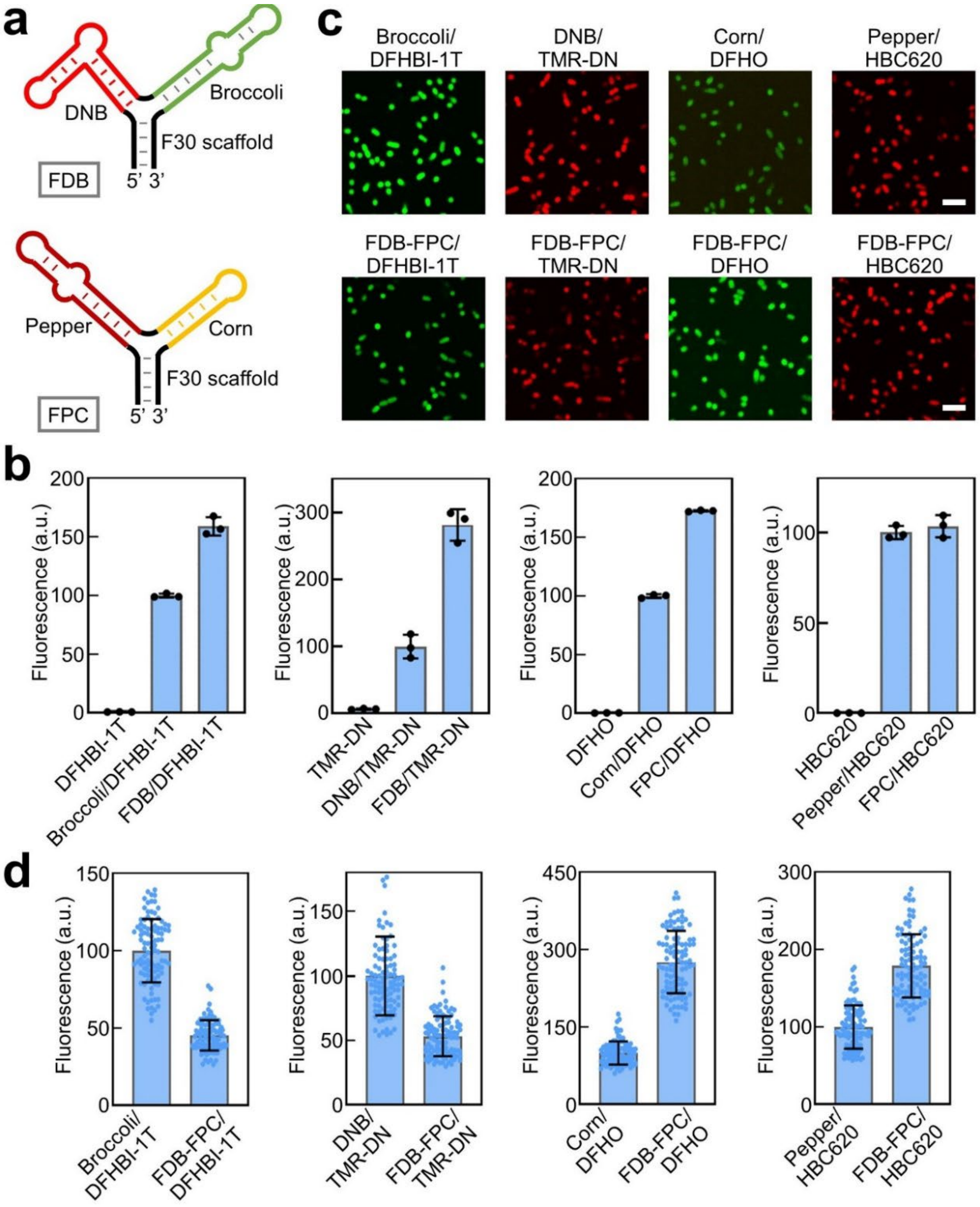


**Fig. S6. Co-expression of four FRs in a pETDuet vector.** (**a**) The constructs of FDB and FPC that contain a DNB/Broccoli- or Pepper/Corn-fused three-way junction F30 RNA scaffold. Each construct was then inserted into a separate expression cassette within the same pETDuet vector. (**b**) Fluorescence intensities as measured in a solution containing 20 μM DFHBI-1T, 0.5 μM TMR-DN, 2 μM DFHO, or 1 μM HBC620 and 1–5 μM corresponding FR, FDB, or FPC. Shown are the mean and the standard error of the mean (SEM) values from three replicated experiments. (**c**) Fluorescence images were taken in BL21 Star (DE3) cells that express Broccoli, DNB, Corn, Pepper, or all four FRs (FDB-FPC) after a 30-min incubation with 200 μM DFHBI-1T, 1 μM TMR-DN, 10 μM DFHO, or 1 μM HBC620 dye molecules. Scale bar, 5 μm. (**d**) Cellular fluorescence intensities as measured in ~50 individual BL21 Star (DE3) cells in each case as shown in the panel (**c**). Shown are the mean and SD values from cellular images taken from at least three independent replicates.

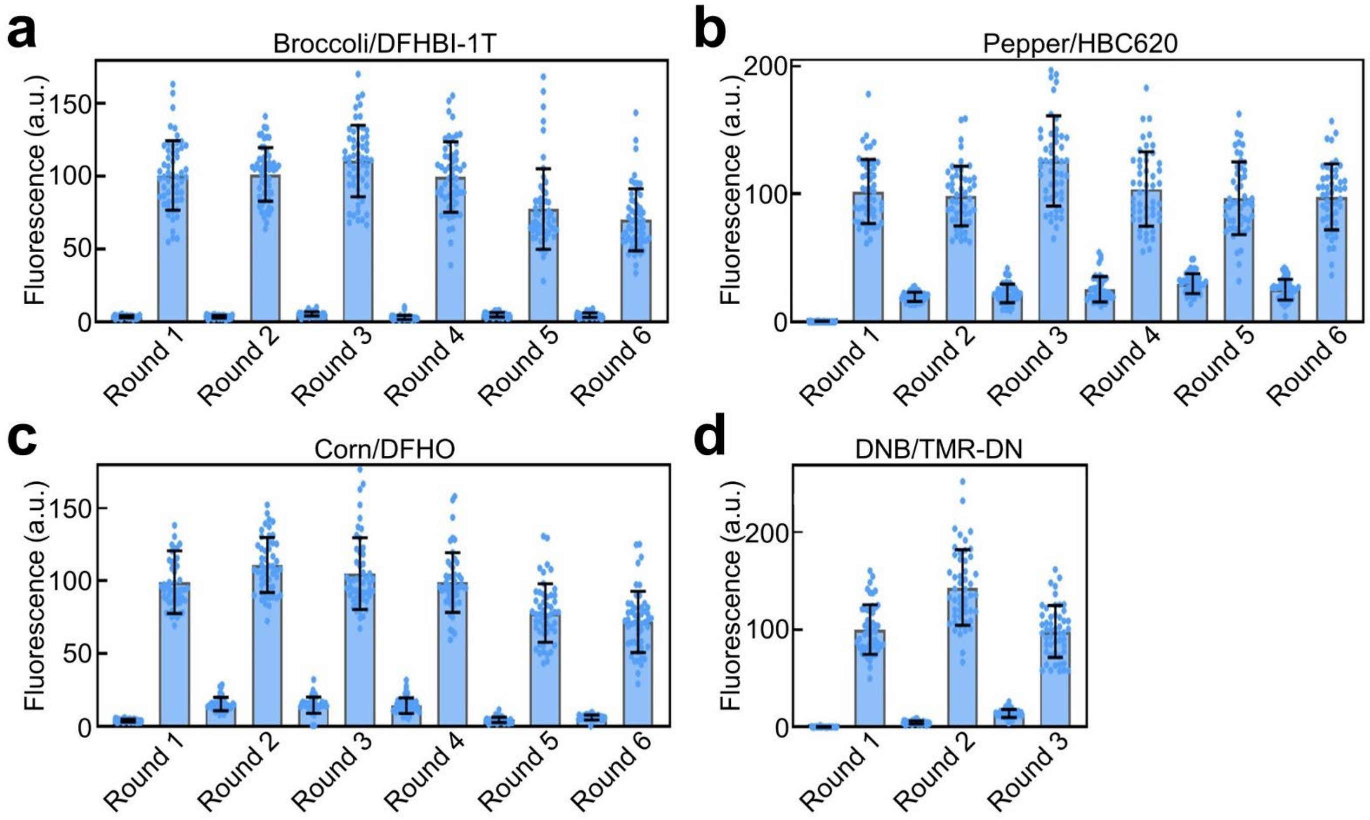


**Fig. S7. Sequential imaging feasibility of FR/dye pairs in live *E. coli* cells.** In BL21 Star (DE3) cells that express all four FRs (FDB-FPC), fluorescence images were taken after each round of 30-min incubation with 200 μM DFHBI-1T, 1 μM TMR-DN, 10 μM DFHO, or 1 μM HBC620. (**a**–**c**) Three times of 1-min DPBS washing in the case of DFHBI-1T, DFHO, and HBC620, or (**d**) five times of 6-min washing for TMR-DN, were performed right after each round of imaging to strip cellular fluorescence. Cellular fluorescence intensities were measured in ~50 individual BL21 Star (DE3) cells in each case. Shown are the mean and SD values from cellular images taken from at least three independent replicates.

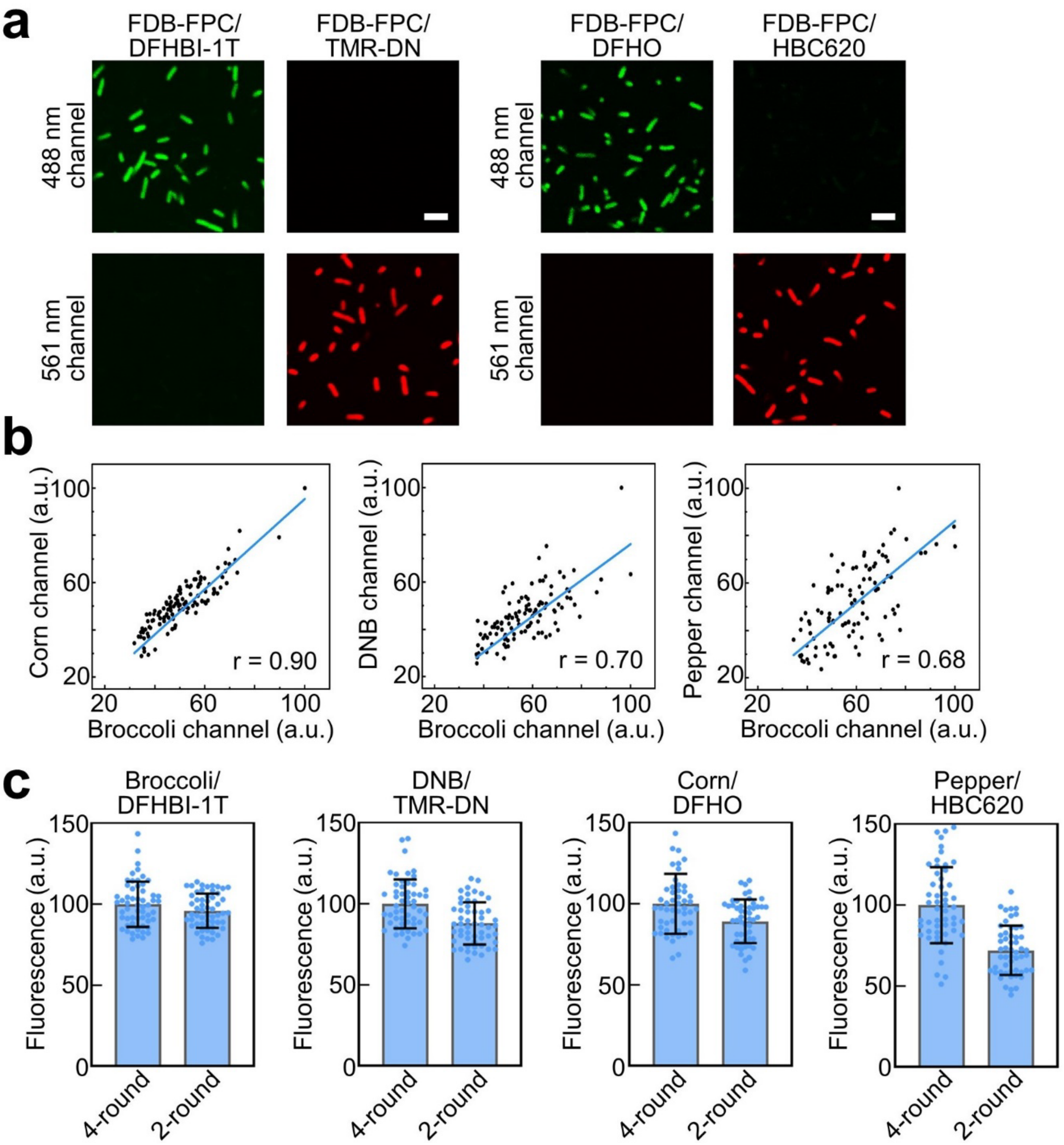


**Fig. S8. Simultaneous detection of two FR/dye pairs to shorten the overall imaging time.** (**a**) Upon excitation via a 488 nm or 561 nm laser, minimal fluorescence interference or bleed-through was observed in BL21 Star (DE3) cells that express all four FRs (FDB-FPC) within a mixture of 200 μM DFHBI-1T and 1 μM TMR-DN, or a mixture of 10 μM DFHO and 1 μM HBC620. Scale bar, 5 μm. (**b**) Correlations of cellular Broccoli, Corn, DNB and Pepper fluorescence signals as measured within ~50 individual cells via a 2-round imaging protocol shown in main Fig. 3c. Pearson's correlation coefficient r was determined in each case where a value between 0.68 and 0.90 indicates a moderate-to-strong positive correlation. (**c**) Cellular fluorescence intensities as measured in ~50 individual BL21 Star (DE3) cells in each case via either a 4-round or 2-round imaging protocol as shown in main Fig. 3. Shown are the mean and SD values from cellular images taken from at least three independent replicates.

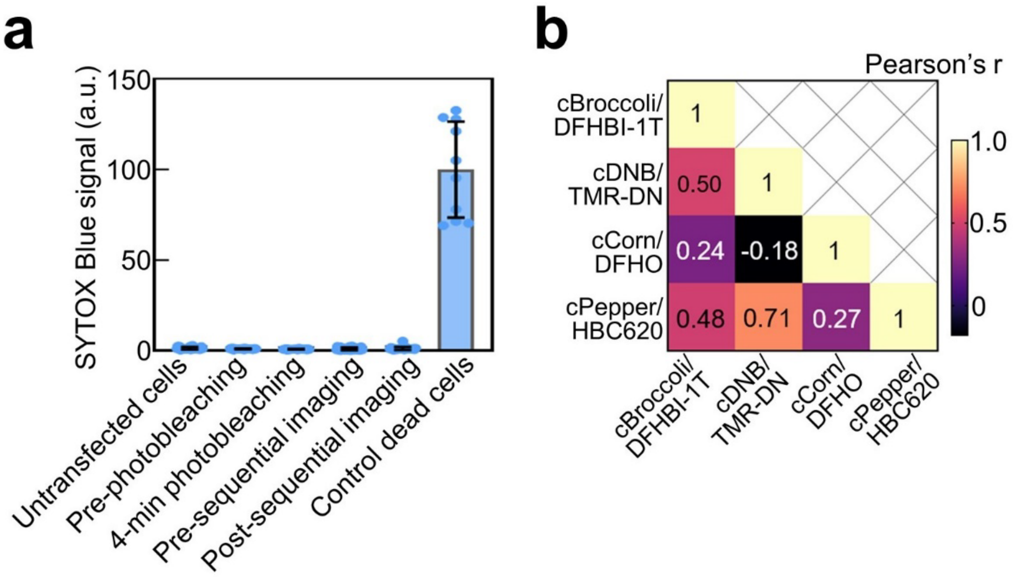


**Fig. S9. Measurement of cytotoxicity and multiplexed imaging correlations in HEK293T cells.** (**a**) Cytotoxicity measurement was performed by incubating HEK293T cells with 1 μM SYTOX Blue for 5 min using untransfected cells (negative control), dead cells (positive cells), and cells before and after a 4-min irradiation with a 1 mW/cm$^2$ power 561 nm laser, as well as cells before and after sequential imaging process shown in main Fig. 5d. Cellular SYTOX Blue fluorescence signals were measured using ~10 individual cells for dead cells and ~30 cells in other cases upon a 405 nm laser irradiation. Shown are the mean and SD values from cellular images taken from at least three independent replicates. (**b**) A correlation heatmap on four FR/dye pair fluorescence signals from 100 individual HEK193T cells as measured via the seqFRIES process shown in main Fig. 5d. The values in the pseudo-color scale showed Pearson's r values as measured between two channels in the x- and y-axis.

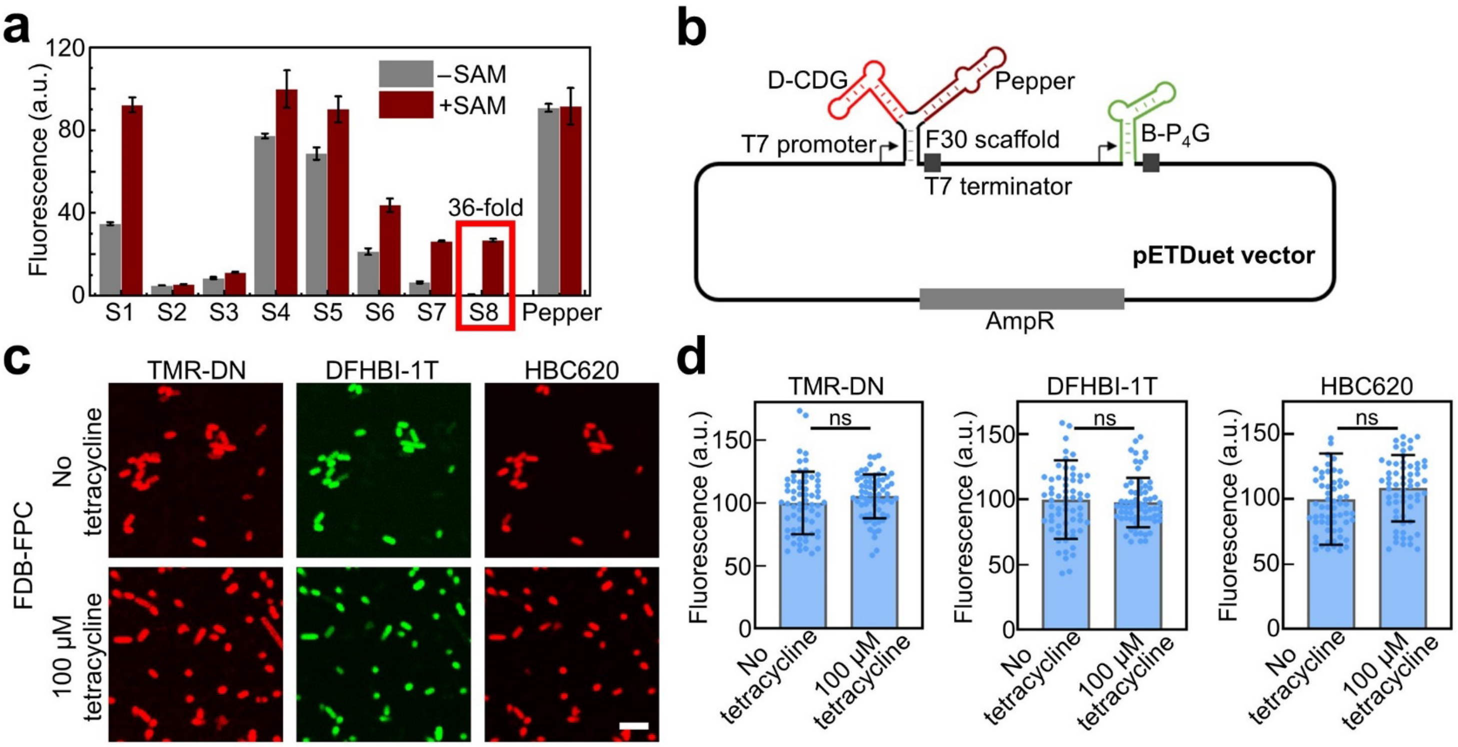


**Fig. S10. seqFRIES-mediated metabolite and signaling molecule sensors.** (**a**) Fluorescence intensities of Pepper-based SAM sensors as measured in a solution containing 1 μM HBC620, 1 μM RNA sensor of different transducer sequences as shown in Table S1, and in the absence or presence of 100 μM SAM. The S8 sensor was identified as the optimal system and named as P-SAM, which exhibited the highest fluorescence enhancement. Shown are the mean and SEM values from at least three independent replicates. (**b**) A pETDuet plasmid construct that co-express FPD-CDG (F30-scaffolded D-CDG sensor and Pepper) and B-$P_4$G. (**c**) Fluorescence images were taken in BL21 Star (DE3) cells that co-express FDB and FPC RNAs. 200 μM DFHBI-1T and 1 μM TMR-DN were added together and incubated for 30 min before the first round of imaging, after three times of 5-min washing, the second round of imaging was performed after a 30-min incubation with 1 μM HBC620. The same seqFRIES procedure was performed in the presence or absence of 100 μM tetracycline. Scale bar, 5 μm. (**d**) The effect of tetracycline on the cellular fluorescence intensities of Broccoli/DFHBI-1T, DNB/TMR-DN, and Pepper/HBC620 was measured in the presence or absence of 100 μM tetracycline. Two-tailed student's t-test: ns, not significant, p>0.05.

### Supplementary Table

**Table S1. List of RNA sequences used in this work.** Blue nucleotides indicate sequences for the F30 scaffold. **Bolded nucleotides** are ribozyme cutting sites for further RNA cyclization. Underlined nucleotides represent the transducer regions for developing FR-based sensors.

| Name | RNA sequence (5' to 3') |
|---|---|
| Broccoli | GGACGGAGACGGUCGGGUCCAGAUAUUCGUAUCUGUCGAGUAGAGUGUGGGCUCCG |
| DNB | GGUGCCUUAUUCCGGACGCCGGGCCCGAAUGCUGCUACGGCAGUCGAAGACAACAUCGCGCCCUUCGGAGGCACC |
| Corn | GGCGCGAGGAAGGAGGUCUGAGGAGGUCACUGCGCC |
| Pepper | GGAUCCCCAAUCGUGGCGUGUCGGCCUGCUUCGGCAGGCACUGGCGCCGGGAUCC |
| Chili | GGCUAGCUGGAGGGGCGCCAGUUCGCUGGUGGUUGGGUGCGGUCGGCUAGCC |
| Mango II | GGAGCACGUACGAAGGAGAGGAGAGGAAGAGGAGAGUACGUGCUCC |
| FDB | GGAAGUUGCCAUGUGUAUCGGGUGCCUUAUUCCGGACGCCGGGCCCGAAUGCUGCUACGGCAGUCGAAGACAACAUCGCGCCCUUCGGAGGCACCCGAUACUCUGAUGAUCCGAGACGGUCGGGUCCAGAUAUUCGUAUCUGUCGAGUAGAGUGUGGGCUCGGAUCAUUCAUGGCAA |
| FPC | GGAAGUUGCCAUGUGUAUCGGGAUCCCCAAUCGUGGCGUGUCGGCCUGCUUCGGCAGGCACUGGCGCCGGGAUCCCGAUACUCUGAUGAUCCGGCGCGAGGAAGGAGGUCUGAGGAGGUCACUGCGCCGGAUCAUUCAUGGCAA |
| cBroccoli | GGCCGCACUCGCCGGUCCCAAGCCCGGAUAAAAUGGGAGGGGGCGGGAAACCGCCU**A**ACCAUGCCGAGUGCGGCCGCUUGCCAUGUGUAUCGGGGAGACGGUCGGGUCCAGAUAUUCGUAUCUGUCGAGUAGAGUGUGGGCUCCCCGAUACUCUGAUGAUCCUUCGGGAUCAUUCAUGGCAAGUGGCCGCGGUCGGCGUGGACUGUA**G**AACACUGCCAAUGCCGGUCCCAAGCCCGGAUAAAAGUGGAGGGUACAGUCCACGC |
| cDNB | GGCCGCACUCGCCGGUCCCAAGCCCGGAUAAAAUGGGAGGGGGCGGGAAACCGCCU**A**ACCAUGCCGAGUGCGGCCGCUUGCCAUGUGUAUCGGGUGCCUUAUUCCGGACGCCGGGCCCGAAUGCUGCUACGGCAGUCGAAGACAACAUCGCGCCCUUCGGAGGCACCCGAUACUCUGAUGAUCCUUCGGGAUCAUUCAUGGCAAGUGGCCGCGGUCGGCGUGGACUGUA**G**AACACUGCCAAUGCCGGUCCCAAGCCCGGAUAAAAGUGGAGGGUACAGUCCACGC |
| cCorn | GGCCGCACUCGCCGGUCCCAAGCCCGGAUAAAAUGGGAGGGGGCGGGAAACCGCCU**A**ACCAUGCCGAGUGCGGCCGCUUGCCAUGUGUAUCGGGCGCGAGGAAGGAGGUCUGAGGAGGUCACUGCGCCCGAUACUCUGAUGAUCCUUCGGGAUCAUUCAUGGCAAGUGGCCGCGGUCGGCGUGGACUGUA**G**AACACUGCCAAUGCCGGUCCCAAGCCCGGAUAAAAGUGGAGGGUACAGUCCACGC |
| cPepper | GGCCGCACUCGCCGGUCCCAAGCCCGGAUAAAAUGGGAGGGGGCGGGAAACCGCCU**A**ACCAUGCCGAGUGCGGCCGCUUGCCAUGUGUAUCGGGAUCCCCAAUCGUGGCGUGUCGGCCUGCUUCGGCAGGCACUGGCGCCGGGAUCCCGAUACUCUGAUGAUCCUUCGGGAUCAUUCAUGGCAAGUGGCCGCGGUCGGCGUGGACUGUA**G**AACACUGCCAAUGCCGGUCCCAAGCCCGGAUAAAAGUGGAGGGUACAGUCCACGC |
| cFDB | GGCCGCACUCGCCGGUCCCAAGCCCGGAUAAAAUGGGAGGGGGCGGGAAACCGCCU**A**ACCAUGCCGAGUGCGGCCGCUUGCCAUGUGUAUCGGGAUCCCCAAUCGUGGCGUGUCGGCCUGCUUCGGCAGGCACUGGCGCCGGGAUCCCGAUACUCUGAUGAUCCGGCGCGAGGAAGGAGGUCUGAGGAGGUCACUGCGC |

| | |
|---|---|
| | CGGAUCAUUCAUGGCAAGUGGCCGCGGUCGGCGUGGACUGUA**G**AACACUG<br>CCAAUGCCGGUCCCAAGCCCGGAUAAAAGUGGAGGGUACAGUCCACGC |
| cFPC | GGCCGCACUCGCCGGUCCCAAGCCCGGAUAAAAUGGGAGGGGGCGGGAAA<br>CCGCCU**A**ACCAUGCCGAGUGCGGCCGCUUGCCAUGUGUAUCGGGAUCCCC<br>AAUCGUGGCGUGUCGGCCUGCUUCGGCAGGCACUGGCGCCGGGAUCCCG<br>AUACUCUGAUGAUCCGGCGCGAGGAAGGAGGUCUGAGGAGGUCACUGCGC<br>CGGAUCAUUCAUGGCAAGUGGCCGCGGUCGGCGUGGACUGUA**G**AACACUG<br>CCAAUGCCGGUCCCAAGCCCGGAUAAAAGUGGAGGGUACAGUCCACGC |
| S1 | GGAUCCCCAAUCGUGGCGUGUCGGCGAAAGGAUGGCGGAAACGCCAGAUG<br>CCUUGUAACCGAAAGGGCACUGGCGCCGGGAUCC |
| S2 | GGAUCCCCAAUCGUGGCGUGUCGCGAAAGGAUGGCGGAAACGCCAGAUGC<br>CUUGUAACCGAAAGGGACUGGCGCCGGGAUCC |
| S3 | GGAUCCCCAAUCGUGGCGUGUCGCCGAAAGGAUGGCGGAAACGCCAGAUG<br>CCUUGUAACCGAAAGGGGACUGGCGCCGGGAUCC |
| S4 | GGAUCCCCAAUCGUGGCGUGUCGUUCCCGAAAGGATGGCGGAAACGCCAG<br>ATGCCTTGTAACCGAAAGGGGGAAACUGGCGCCGGGAUCC |
| S5 | GGAUCCCCAAUCGUGGCGUGUCGAAGCGAAAGGATGGCGGAAACGCCAGA<br>TGCCTTGTAACCGAAAGGGCUUACUGGCGCCGGGAUCC |
| S6 | GGAUCCCCAAUCGUGGCGUGUCGGGCGAAAGGAUGGCGGAAACGCCAGAU<br>GCCUUGUAACCGAAAGGGCCACUGGCGCCGGGAUCC |
| S7 | GGAUCCCCAAUCGUGGCGUGUCGUUCGAAAGGATGGCGGAAACGCCAGAT<br>GCCTTGTAACCGAAAGGGAAACUGGCGCCGGGAUCC |
| S8 (P-SAM) | GGAUCCCCAAUCGUGGCGUGUCGAAGAAAGGATGGCGGAAACGCCAGATG<br>CCTTGTAACCGAAAGGUUACUGGCGCCGGGAUCC |
| B-$P_4$G | GGAAGUGUACCUUAGGGUUCCGGCCAUAAGGCGUCAGCGACCGAGCGG<br>UACAAUGCUGUCGAGUAGAGUGUGGGCUCGCAAGAGACGGUCGGGUCCA<br>GCAACACCGUGAGCAUAAAAGGCUCCAGCGGCAAGUUCC |
| D-CDG | GGUGCCUUAUUCCGGACGCCGGGCCCGAAUCUUCGAUAACGGCAAACUUG<br>UCGAAAGAUAAGGACGCAAAGCCACAGGGCCUUCUUGAUGAACCGUCAAUA<br>GGCAGCCUGGCUACCGAAGCGAAGACAACAUCGCGCCCUUCGGAGGCACC |
| C-SAH | GGGCUCGCCGAGGAGCGCUGCAAGGAAGAGGAAGGAGGUCUGAGGAGGU<br>CACUUUCCCCAGGCUCGGCGAUUAUCGGGACCUUUAAACCAACGGCGCUC<br>GGUUCAG |
